# Monitoring the microbiome for food safety and quality using deep shotgun sequencing

**DOI:** 10.1101/2020.05.18.102574

**Authors:** Kristen L. Beck, Niina Haiminen, David Chambliss, Stefan Edlund, Mark Kunitomi, B. Carol Huang, Nguyet Kong, Balasubramanian Ganesan, Robert Baker, Peter Markwell, Ban Kawas, Matthew Davis, Robert J. Prill, Harsha Krishnareddy, Ed Seabolt, Carl H. Marlowe, Sophie Pierre, André Quintanar, Laxmi Parida, Geraud Dubois, James Kaufman, Bart C. Weimer

## Abstract

In this work, we hypothesized that shifts in the food microbiome can be used as an indicator of unexpected contaminants or environmental changes. To test this hypothesis, we sequenced total RNA of 31 high protein powder (HPP) samples of poultry meal pet food ingredients. We developed a microbiome analysis pipeline employing a key eukaryotic matrix filtering step that improved microbe detection specificity to >99.96% during *in silico* validation. The pipeline identified 119 microbial genera per HPP sample on average with 65 genera present in all samples. The most abundant of these were *Bacteroides, Clostridium, Lactococcus, Aeromonas*, and *Citrobacter.* We also observed shifts in the microbial community corresponding to ingredient composition differences. When comparing culture-based results for *Salmonella* with total RNA sequencing, we found that *Salmonella* growth did not correlate with multiple sequence analyses. We conclude that microbiome sequencing is useful to characterize complex food microbial communities, while additional work is required for predicting specific species’ viability from total RNA sequencing.

## 1. INTRODUCTION

Sequencing the microbiome of food may reveal characteristics about the associated microbial content that culturing or targeted whole genome sequencing alone cannot. However, to meet the various needs of food safety and quality, next generation sequencing (NGS) and analysis techniques require additional development^1^ with specific consideration for accuracy, speed, and applicability across the supply chain.^2^ Microbial communities and their characteristics have been studied in relation to flavor and quality in fermented foods,^3–5^ agricultural processes in grape^6^ and apple fruit^7^, and manufacturing processes and production batches in Cheddar cheese.^8^ However, the advantage of using the microbiome specifically for food safety and quality has yet to be demonstrated.

Currently, food safety regulatory agencies including the Food and Drug Administration (FDA), Centers for Disease Control and Prevention (CDC), United States Department of Agriculture (USDA), and European Food Safety Authority (EFSA) are converging on the use of whole genome sequencing (WGS) for pathogen detection and outbreak investigation. Large scale WGS of food-associated bacteria was first initiated via the 100K Pathogen Genome Project^9^ with the goal of expanding the diversity of bacterial reference genomes— a crucial need for foodborne illness outbreak investigation, traceability, and microbiome studies.^10,11^ However, since WGS relies on culturing a microbial isolate prior to sequencing, there are inherent biases and limitations in its ability to describe the microorganisms and their interactions in a food sample. Such information would be very valuable for food safety and quality applications.

High throughput sequencing of total DNA and total RNA are promising approaches to characterize microbial niches in their native state without introducing bias due to culturing.^12–14^ Additionally, total RNA sequencing has the potential to provide evidence of live and biologically active components of the sample.^14,15^ It also provides accurate microbial naming, relative microbial abundance, and better reproducibility than total DNA or amplicon sequencing.^14^ Total RNA sequencing minimizes PCR amplification bias that occurs in single gene amplicon sequencing and overcomes the decreased detection sensitivity from using DNA sequencing in metagenomics.^14^ Total RNA metatranscriptome sequencing, however, is yet to be examined in raw food ingredients as a method to provide robust characterization of the microbial communities and the interacting population dynamics.

From a single sequenced food microbiome, numerous dimensions of the sample can be characterized that may yield important indicators of safety and quality. Using total DNA or RNA, evidence for the eukaryotic food matrix can be examined. In Haiminen *et al*.,^16^ we quantitatively demonstrated the utility of metagenome sequencing to authenticate the composition of complex food matrices. In addition, from total DNA or RNA, one can observe signatures from commensal microbes, pathogenic microbes, and genetic information for functional potential (from DNA) or biologically active function (from RNA).^14,15^ Detecting active transcription from live microbes in food is very important to avoid spurious microbial observations that may instead be false positives due to quiescent DNA in the sample. Use of RNA in food analytics also offers the opportunity to examine expression of metabolic processes that are related to antibiotic resistance,^17,18^ virulence factors, or replication genes, among others. Additionally, it has the potential to define viable microbes that are capable of replication in the food and even microorganisms that stop replicating but continue to produce metabolic activity that changes food quality and safety.^19–24^

Microorganisms are sensitive to changes in temperature, salinity, pH, oxygen content, and many other physicochemical factors that alter their ability to grow, persist, and cause disease. They exist in dynamic communities that change in response to environmental perturbation – just as the gut microbiome shifts in response to diet.^25–28^ Shifts in microbiome composition or activity can be leveraged in the application of microbiome characterization to monitor the food supply chain. For example, Noyes et al. followed the microbiome of cattle from the feed lot to the food packaging, concluding that the microbial community and antibiotic resistance characteristics change based on the processing stage.^17,18,29^ We hypothesize that observable shifts in microbial communities of food can serve as an indicator of food quality and safety.

In this work, we examined 31 high protein powder samples (HPP; derived from poultry meal). HPP are commonly used raw materials in pet foods. They are subject to microbial growth prior to preparation and continued survival in powder form.^30^ We subjected the HPP samples to deep total RNA sequencing with ∼300 million reads per sample. In order to process the 31 samples collected over ∼1.5 years from two suppliers at a single location, we defined and calibrated the appropriate methods– from sample preparation to bioinformatic analysis– needed to taxonomically identify the community members present and to detect key features of microbial growth. First, we removed the HPP’s food matrix RNA content as eukaryotic background with an important bioinformatic filtering step designed specifically for food analysis. The remaining sequences were used for relative quantification of microbiome members and for identifying shifts based on food matrix content, production source, and *Salmonella* culturability. This work demonstrates that total RNA sequencing is a robust approach for monitoring the food microbiome for use in food safety and quality applications, while additional work is required for predicting pathogen viability.

## 2. RESULTS

### 2.1 Evaluation of microbial identification capability in total RNA and DNA sequencing

Microbial identification in microbiomes often leverages shotgun DNA sequencing; however, total RNA sequencing can provide additional information about viable bacterial activity in a community via transcriptional activity. Since using total RNA to study food microbiomes is novel, each step of the analysis workflow (Figure 1) was carefully designed and scrutinized for accuracy. For all analyses done in this study, we report relative abundance in reads per million (RPM) (Equation 1) as recommended by Gloor et al^31,32^ and apply the conservative threshold of RPM > 0.1 to indicate presence as previously described by Langelier et al and Illot et al.^33,34^ Numerically, this threshold translates to ∼30 reads per genus per sample considering a sequencing depth of ∼300 million reads per sample (Methods Section 4.4). First, we examined the effectiveness of RNA for taxonomic identification and relative quantification of microbes in the presence of food matrix reads. We observed that RNA sequencing results correlated (R^2^ = 0.93) with the genus relative quantification provided by DNA sequencing (Supplementary Figure S1). RNA sequencing also detected more genera demonstrated by a higher *α*-diversity than the use of DNA (Supplementary Figure S2). Additionally, from the same starting material, total RNA sequencing resulted in 2.4-fold more reads classified to microbial genera compared to total DNA sequencing (after normalizing for sequencing depth). This increase is substantial as microbial reads are such a small fraction of the total sequenced reads. Considering these results, we further examined the microbial content from total RNA extracted from 31 high protein powder (HPP) samples (Supplementary Table 1) that resulted in an average of ∼300 million paired end 150 bp sequencing reads per sample in this study.

**Figure 1:**
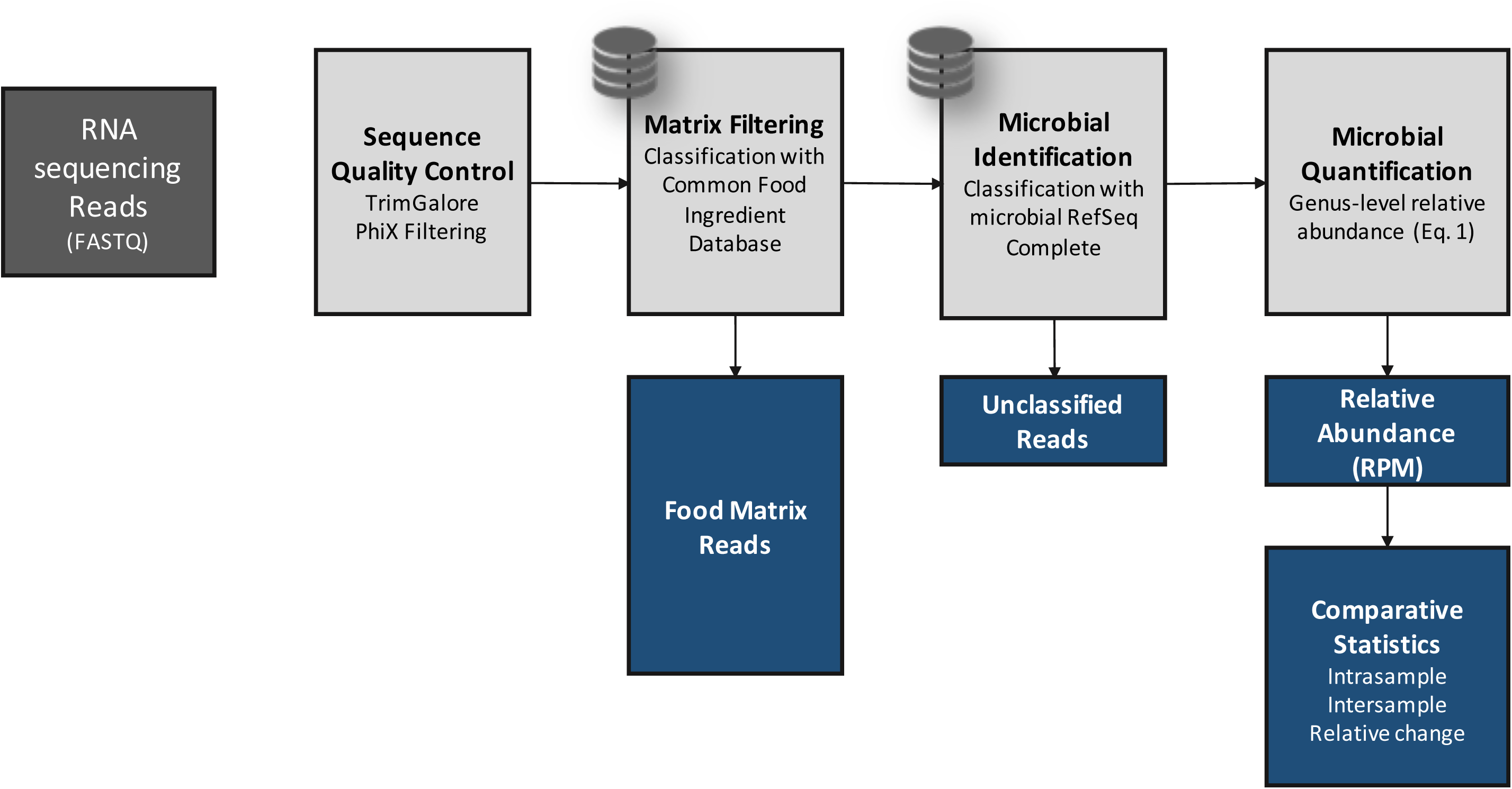
Pipeline description of bioinformatic steps applied to high protein powder metatranscriptome samples. Black arrows indicate data flow and blue boxes describe outputs from the pipeline.

### 2.2 Evaluation and application of *in silico* filtering of eukaryotic food matrix reads

Sequenced reads from the eukaryotic host or food matrix may lead to false positives for microbial identification in microbiome studies.^35^ This may occur partly due to reads originating from low complexity regions of eukaryotic genomes, e.g. telomeric and centromeric repeats, being misclassified as spurious microbial hits.^36^ In total DNA or RNA sequencing of clinical or animal or even plant microbiomes, eukaryotic content may often comprise > 90% of the total sequencing reads. This presents an important bioinformatic challenge that we addressed by filtering matrix content using a custom-built reference database of 31 common food ingredient and contaminant genomes (Supplementary Table 2) using the *k*-mer classification tool Kraken.^37^ This step allows for rapidly classifying all sequenced reads (∼300 million reads for each of 31 samples) as matrix or non-matrix. The matrix filtering process yielded an estimate of the total percent matrix content for a sample. See our work in Haiminen et al.^38^ on quantifying the eukaryotic food matrix components with further precision.

To validate the matrix filtering step, we constructed *in silico* mock food microbiomes with a high proportion of complex food matrix content and low microbial content (Supplementary Table 3). We then computed the true positive, false positive, and false negative rates of observed microbial genera and sequenced reads (Table 1). False positive viral, archaeal, and eukaryotic microbial genera (as well as bacteria) were observed without matrix filtering, although bacteria were the only microbes included in the simulated mixtures. Introducing a matrix filtering step to the pipeline improved read classification specificity to >99.96% (from 78–93% without filtering) in both simulated food mixtures, while maintaining zero false negatives. With this level of demonstrated accuracy, we used bioinformatic matrix filtering prior to further microbiome analysis.

**Table 1:** Microbial Identification Accuracy from Simulated Food Microbiome Mixtures. Accuracy of microbial identification using *in silico* constructed Simulated Food Mixtures with expected food matrix and microbial sequences.

| Table 1: Microbial Identification Accuracy from Simulated Food Microbiome Mixtures |  |  |  |  |  |  |  |  |  |
| --- | --- | --- | --- | --- | --- | --- | --- | --- | --- |
|  | Simulated Mixture 1 |  |  |  |  | Simulated Mixture 2 |  |  |  |
|  | With Matrix Filtering |  | No Matrix Filtering |  |  | With Matrix Filtering |  | No Matrix Filtering |  |
|  | # GENERA | GENUS READS | # GENERA | GENUS READS |  | # GENERA | GENUS READS | # GENERA | GENUS READS |
| Bacteria in Simulated Mixture (Expected Content) | 14 | 15,000 | 14 | 15,000 |  | 14 | 15,000 | 14 | 15,000 |
| Observed Microbial Content |  |  |  |  |  |  |  |  |  |
| Bacteria | 18 | 13,517 | 34 | 13,700 |  | 15 | 13,551 | 33 | 13,999 |
| Viruses | 0 | 0 | 9 | 563 |  | 0 | 0 | 4 | 328 |
| Archaea | 0 | 0 | 1 | 1 |  | 0 | 0 | 1 | 3 |
| Eukaryota | 0 | 0 | 4 | 104 |  | 0 | 0 | 4 | 799 |
| Total Observed | 18 | 13,517 | 48 | 14,368 |  | 15 | 13,551 | 42 | 15,129 |
| True Positives<br>(as a % of total observed) | 14<br>(78%) | 13,511<br>(99.96%) | 14<br>(29%) | 13,571<br>(94.45%) |  | 14<br>(93%) | 13,548<br>(99.98%) | 14<br>(33%) | 13,623<br>(90.05%) |
| False Positives<br>(as a % of total observed) | 4<br>(22%) | 6<br>(0.04%) | 34<br>(71%) | 797<br>(5.55%) |  | 1<br>(7%) | 3<br>(0.02%) | 28<br>(67%) | 1,506<br>(9.95%) |
| False Positives Removed with Matrix Filtering<br>(as a % of false positives without filtering) | 30<br>(88.2%) | 791<br>(99.2%) |  |  |  | 27<br>(96.4%) | 1,503<br>(99.8%) |  |  |

### 2.3 High protein powder microbiome ecology

After filtering eukaryotic matrix sequences, we applied the remaining steps in the bioinformatic workflow (Figure 1) to examine the shift in the high protein powder (HPP) microbiome membership and to quantify the relative abundance of microbes at the genus level. Genus is the first informative taxonomic rank for food pathogen identification that can be considered accurate given current incompleteness of reference databases^11,39–42^ and was therefore used in subsequent analyses. Overall, between 98 and 195 microbial genera (avg. 119) were identified (RPM > 0.1) per HPP sample (Supplementary Table 4). When analyzing *α*-diversity i.e. the number of microbes detected per sample, inter-sample comparisons may become skewed unless a common number of reads is considered since deeper sequenced samples may contain more observed genera merely due to a greater sampling depth.^43,44^ Thus, we utilized bioinformatic rarefaction i.e. subsampling analysis to showcase how microbial diversity was altered by sequencing depth. Examination of *α*-diversity across a range of *in silico* subsampled sequencing depths showed that the community diversity varied across samples (Figure 2A). One sample (MFMB-04) had 1.7 times more genera (195) than the average across other samples (avg. 116, range 98–143) and exhibited higher *α*-diversity than any other sample at each *in silico* sampled sequencing depth (Figure 2A). Rarefaction analysis further demonstrated that when considering fewer than ∼67 million sequenced reads, the observable microbial population was not saturated (median elbow calculated as indicated in Satopää, et al.^45^). This observation suggests that deeper sequencing or more selective sequencing of the HPP microbiomes will reveal more microbial diversity.

**Figure 2A:**
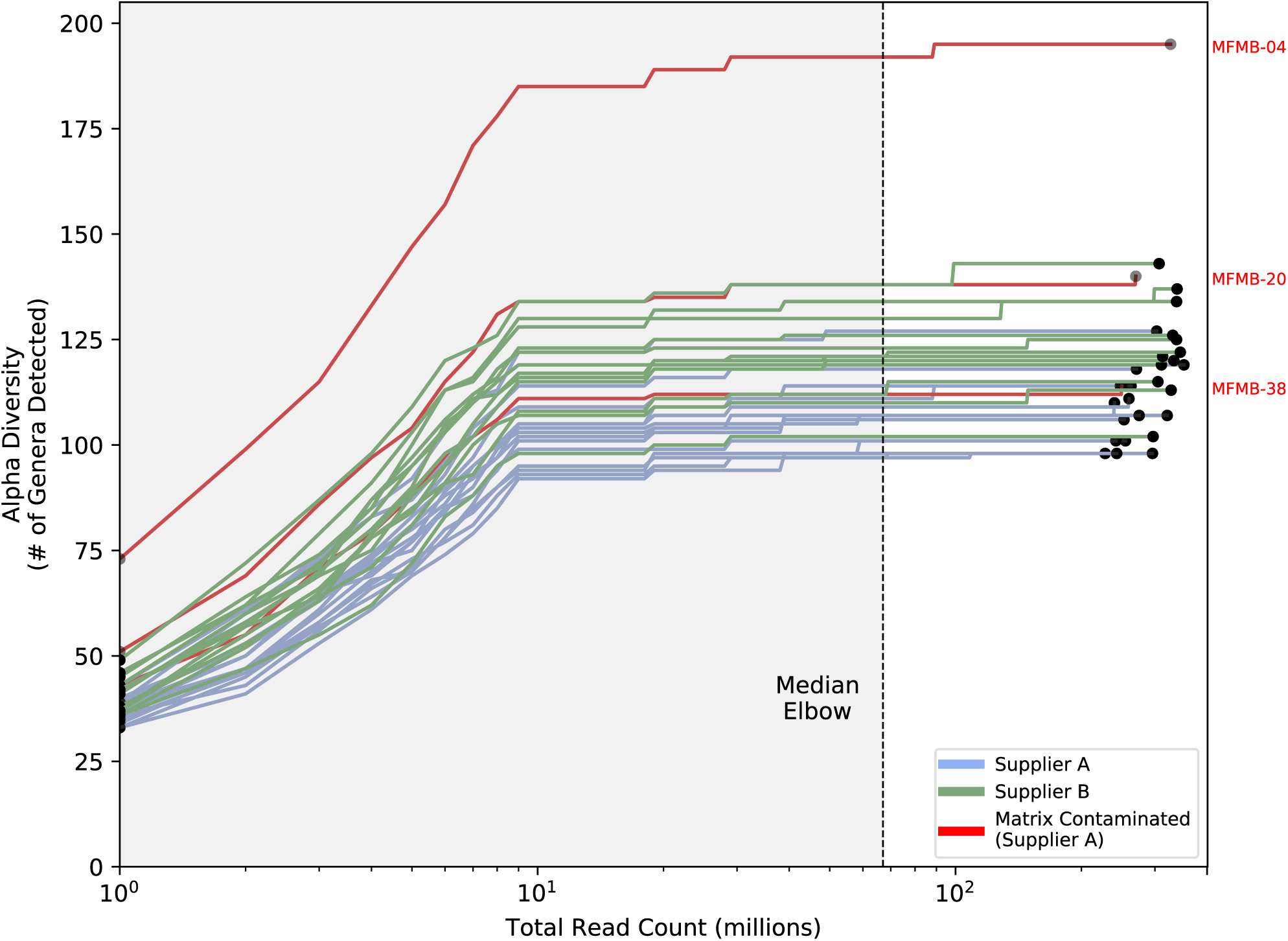
Alpha diversity (number of genera) for all (n = 31) high protein powder metatranscriptomes is compared to total number of sequenced reads for a range of *in silico* subsampled sequencing depths. The dashed vertical line indicates the median elbow (at approx. 67 million reads).

Notably, between 2%–4% (approximately 5,000,000–14,000,000) of reads per sample remained unclassified as either eukaryotic matrix or microbe (Supplementary Figure S3). However, the unclassified reads exhibited a GC (guanine plus cytosine) distribution similar to reads classified as microbial (Supplementary Figure S4) indicating these reads may represent microbial content that is absent or sufficiently divergent from existing references.

We calculated *β*-diversity to study inter-sample microbiome differences and to identify any potential outliers among the sample collection. The Aitchison distances^46^ of microbial relative abundances were calculated between samples (as recommended for compositional microbiome data^31,32^), and the samples were hierarchically clustered based on the resulting distances (Figure 2B). The two primary clades were mostly defined by supplier (except for MFMB-17). In Haiminen *et al*.,^38^ we reported that three of the HPP samples contained unexpected eukaryotic species. We hypothesized that the presence of these contaminating matrix components (beef identifiable as *Bos taurus* and pork identifiable as *Sus scrofa*) would alter the microbiome as compared to chicken (identifiable as *Gallus gallus*) alone. Clustering HPP samples using their microbiome membership led to a distinctly different group of the matrix-contaminated samples, supporting this hypothesis (Figure 2B). These observations indicate that samples can be discriminated based on their microbiome content for originating source and supplier, which is necessary for source tracking potential hazards in food.

**Figure 2B:**
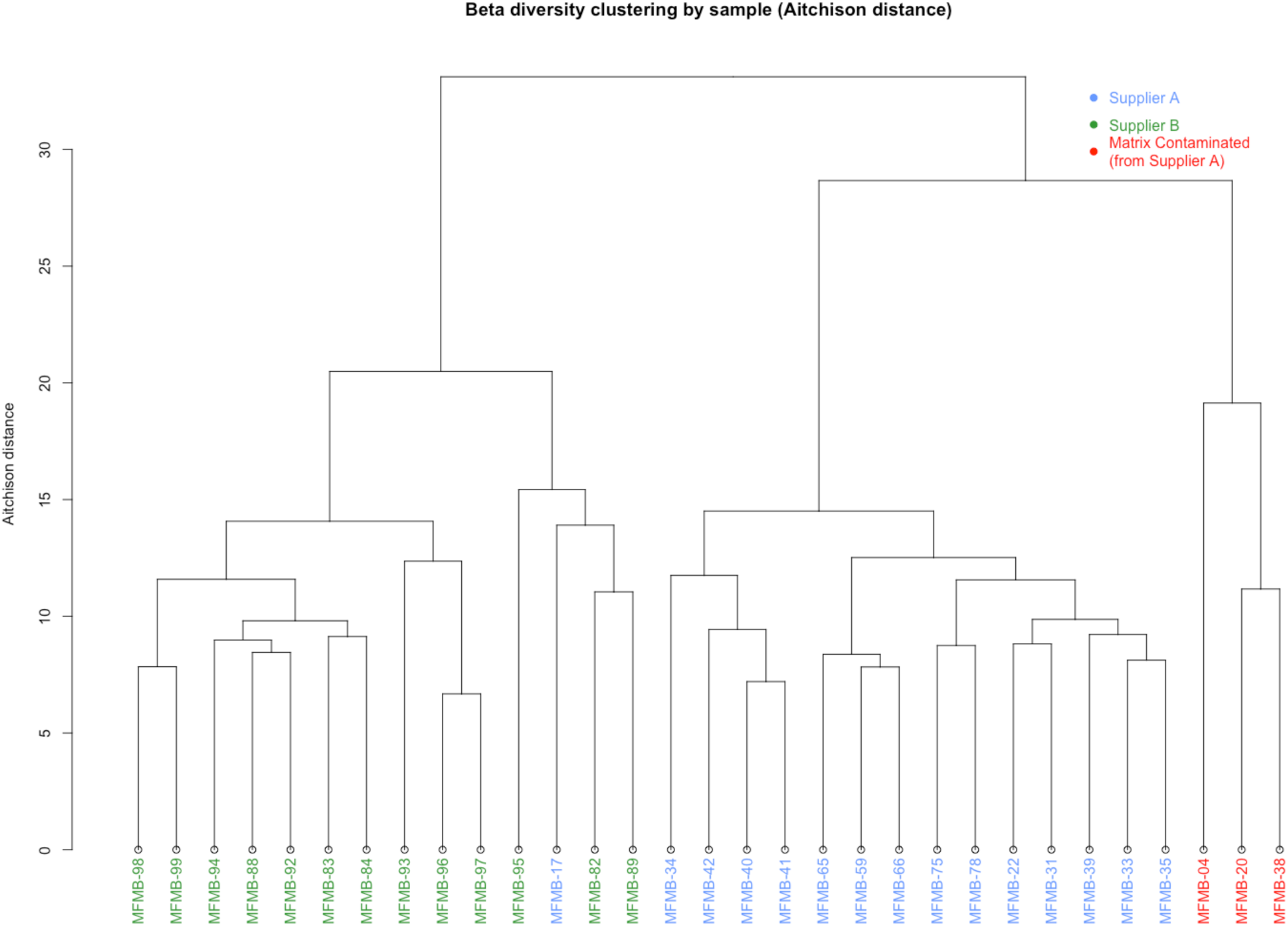
Hierarchical clustering of Aitchison distance values of poultry meal samples based on microbial composition. Samples were received from Supplier A (blue and red) and Supplier B (green). Matrix-contaminated samples are additionally marked in red.

### 2.4 Comparative analysis of high protein powder microbiome membership and composition

We identified 65 genera present in all HPP samples (Figure 3A), whose combined abundance accounted for between 88-99% of the total abundances of detected genera per sample. *Bacteroides, Clostridium, Lactococcus, Aeromonas*, and *Citrobacter* were the five most abundant of these microbial genera. The identified microbial genera also included viruses, the most abundant of which was *Gyrovirus* (< 10 RPM per sample). *Gyrovirus* represents a genus of non-enveloped DNA viruses responsible for chicken anemia which is ubiquitous in poultry. While there were only 65 microbial genera identified in all 31 HPP samples, the *α*-diversity per sample was on average two-fold greater as previously indicated.

**Figure 3A:**
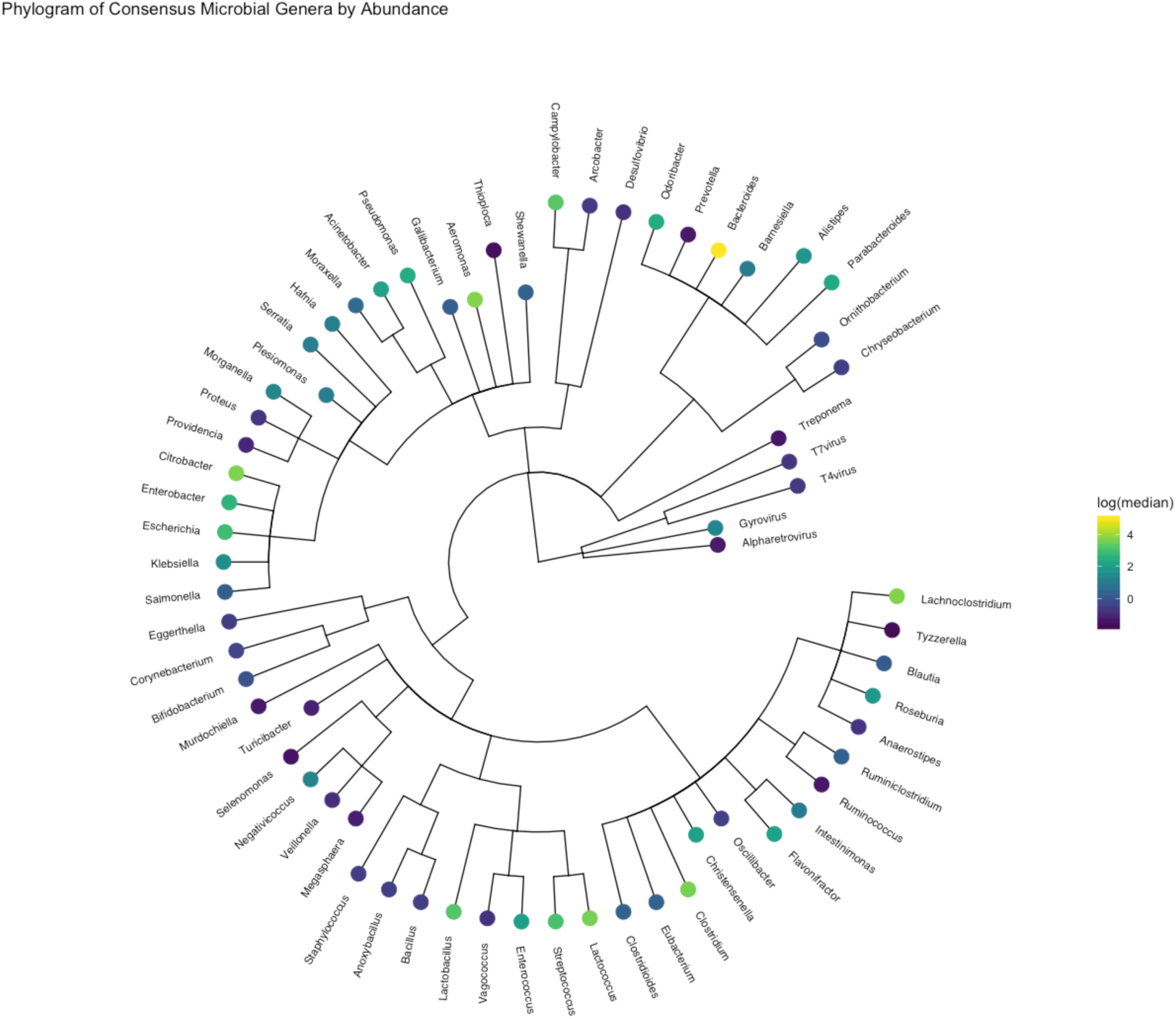
Phylogram of the 65 microbial genera present in all samples with RPM > 0.1

Beyond the collection of 65 microbes observed in all samples, there were an additional 164 microbes present in various HPP samples. Together, we identified a total of 229 genera among the 31 HPP samples tested (Figure 3B and 4, Supplementary Table 4). In order to identify genera that were most variable between samples, we computed the median absolute deviation (MAD)^47^ using the normalized relative abundance of each microbe (Figure 5A). The abundance of *Bacteroides* was the most variable among samples (median = 148.1 RPM, MAD = 30.6) and showed increased abundance in almost all samples from Supplier A compared to Supplier B (abundance for the 10 most variable genera shown in Figure 5B). *Clostridium* (median = 37.4 RPM, MAD = 24.2), *Lactococcus* (median = 36.8 RPM, MAD = 18.2), and *Lactobacillus* (median = 24.2, MAD = 7.2) were also highly variable and 3–4 fold more abundant in samples MFMB-04 and MFMB-20 compared to other samples (Figure 5B). *Pseudomonas* (median = 11.1 RPM, MAD = 12.2) was markedly more abundant in MFMB-83 than any other sample (Figure 5B). These genera highlight variability between microbiomes from a single food source and may provide insights into other dissimilarities in these samples.

**Figure 3B:**
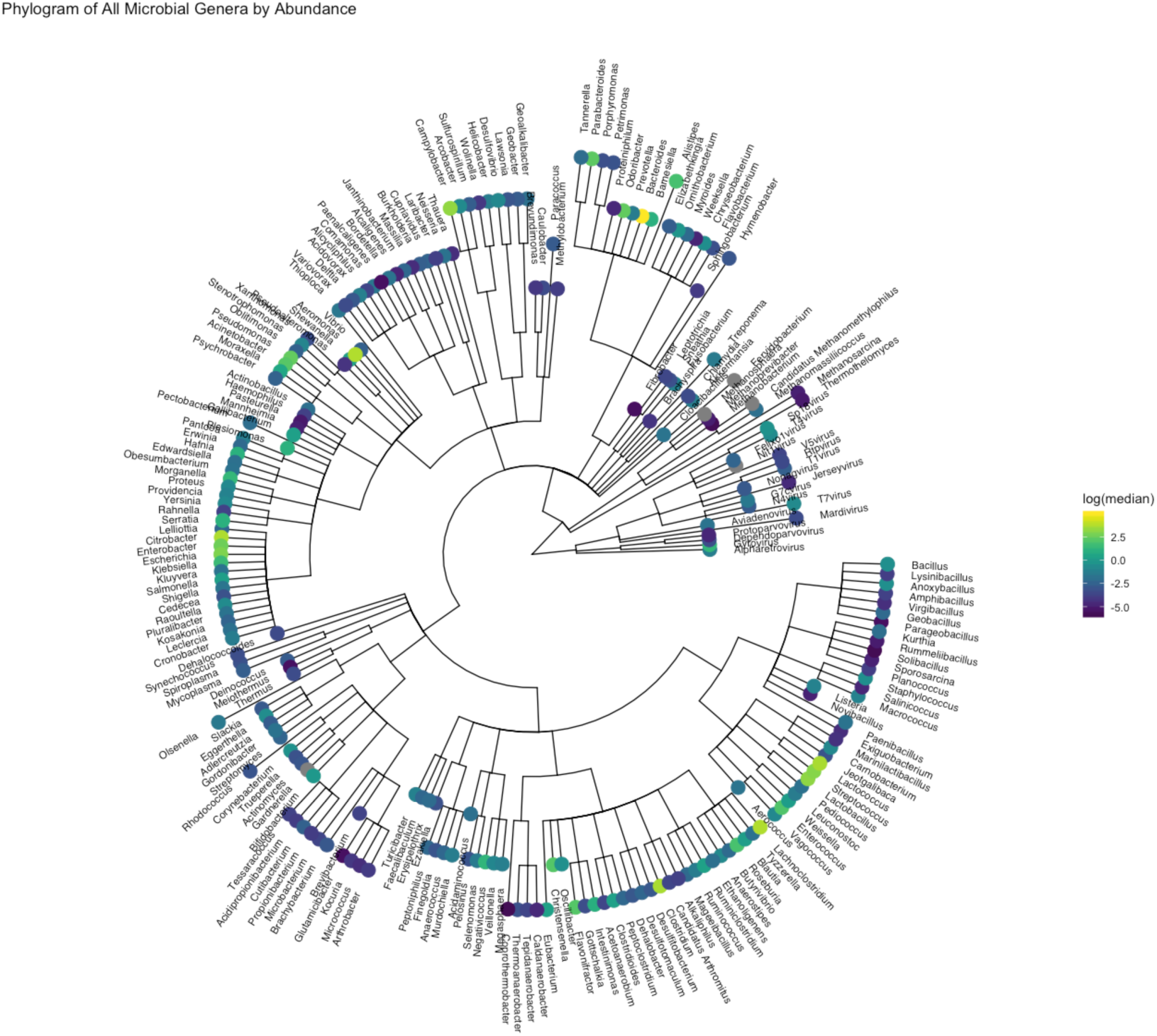
Phylogram of all microbes observed in *any* sample. Log of the median RPM value across samples is indicated. Grey indicating a median RPM value of 0.

**Figure 4:**
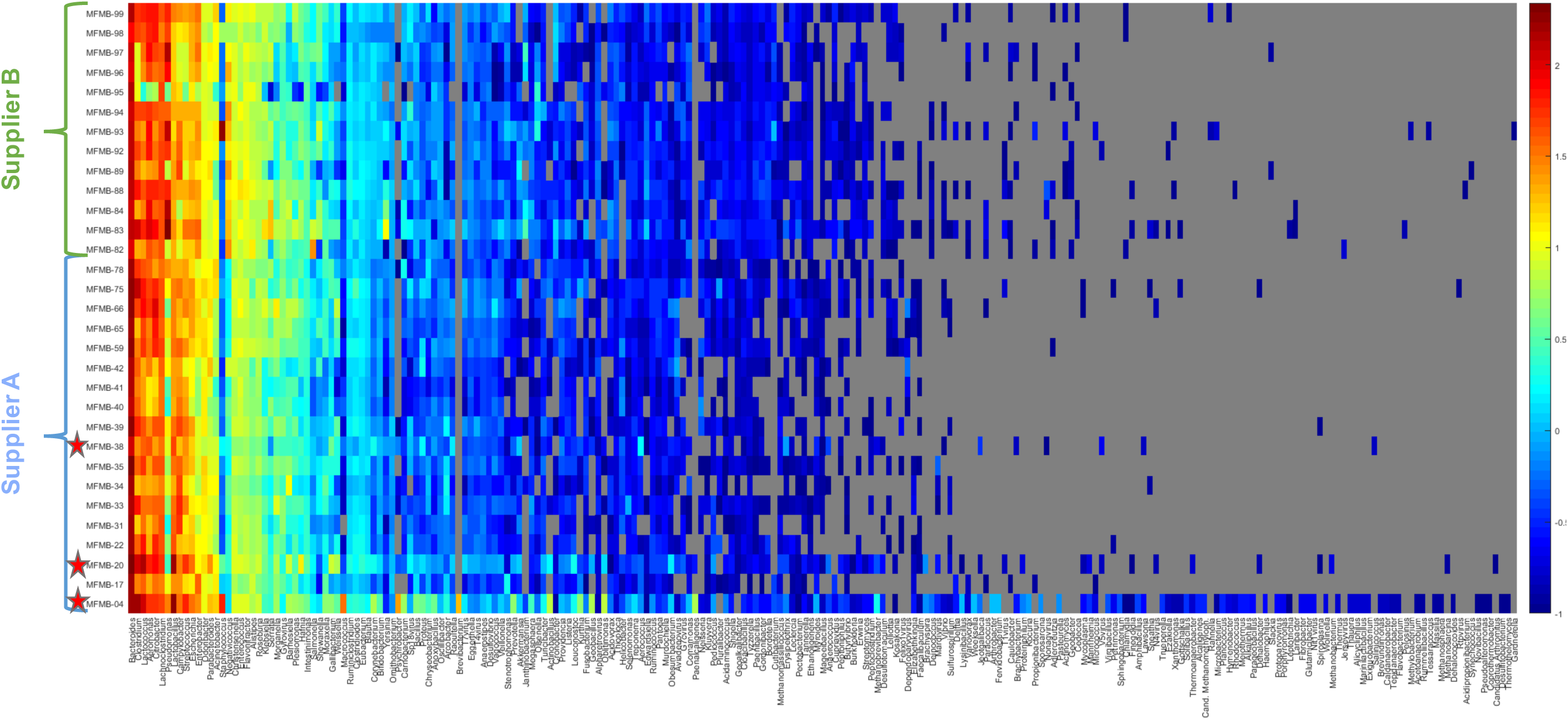
Heatmap (log_10_-scale) of high protein powder microbial composition and relative abundance (RPM) where absence (RPM < 0.1) is indicated in grey. Genera are ordered by summed abundance across samples. Samples were received from Supplier A (blue) and Supplier B (green). Red stars indicate matrix-contaminated samples (from Supplier A).

**Figure 5A:**
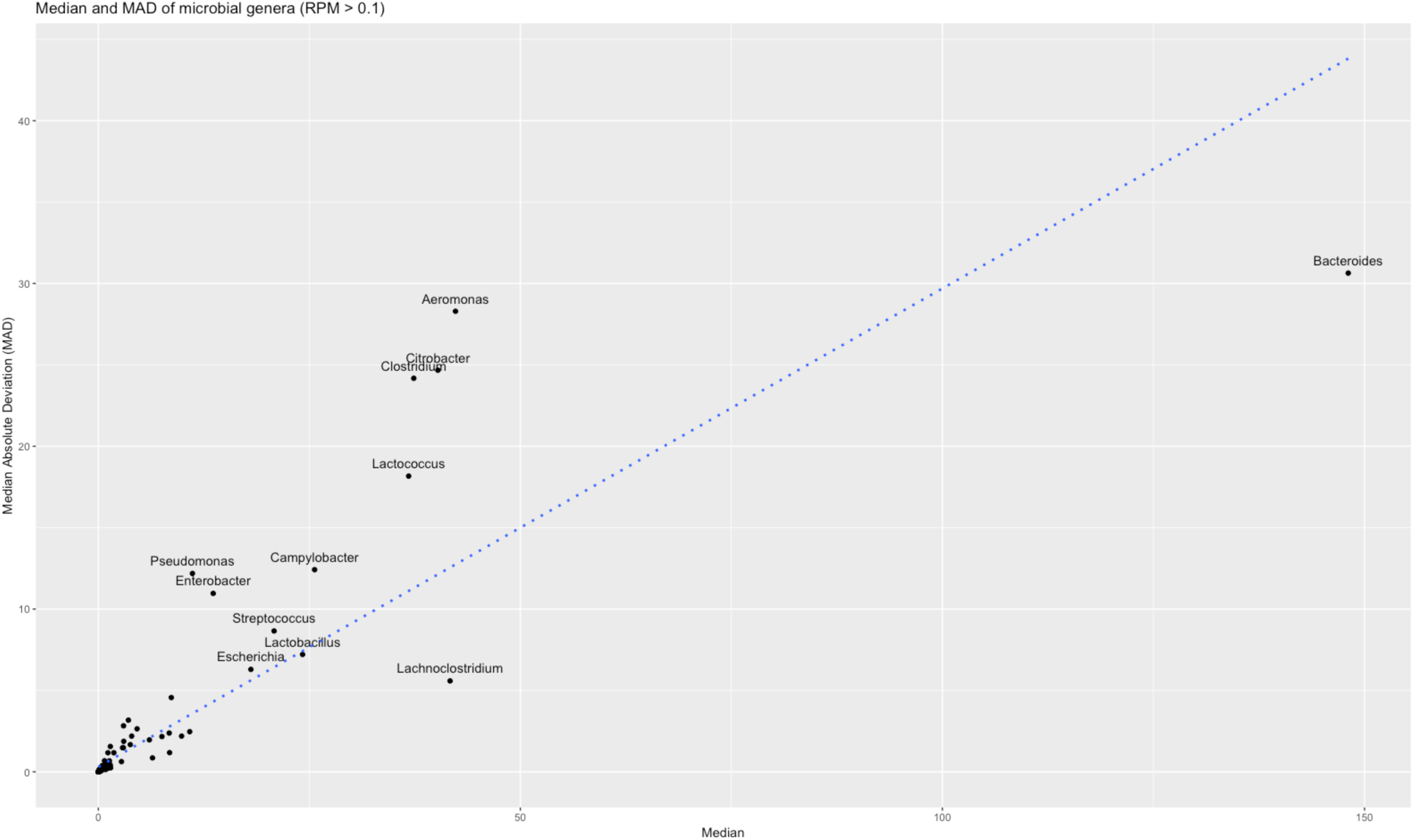
All identified microbial general are plotted with median value and median absolute deviation (MAD) of RPM abundance. Genera with MAD > 5 are labeled with the genus name.

**Figure 5B:**
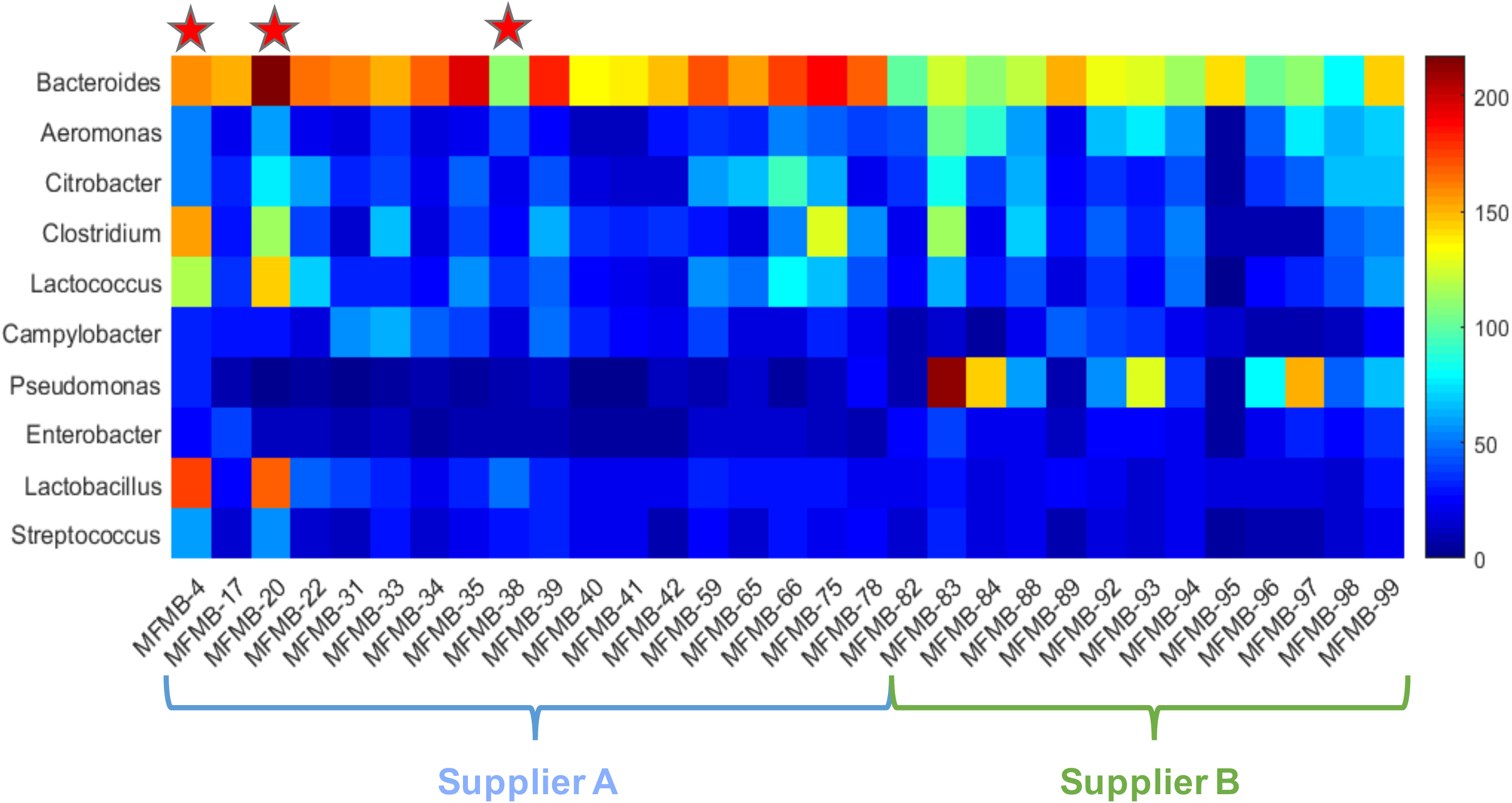
Heatmap (log_10_-scale) of ten microbial genera with the largest median absolute deviation (MAD) across samples. Genera are ordered by decreasing MAD from top to bottom. Samples were received from Supplier A (blue) and Supplier B (green). Red stars indicate matrix contaminated samples (from Supplier A).

### 2.5. Microbiome shifts in response to changes in food matrix composition

We tested the hypothesis that the microbiome composition will shift in response to changes in the food matrix and can be a unique signal to indicate contamination or adulteration. In 28 of the 31 HPP samples, >99% of the matrix reads were determined in our related work^38^ to originate from poultry (*Gallus gallus*), which was the only ingredient expected based on ingredient specifications. However, three samples had higher pork and beef content compared to all other HPP samples: MFMB-04 (7.74% pork, 8.99% beef), MFMB-20 (0.53% pork, 1.00% beef), and MFMB-38 (0.92% pork, 0.29% beef) compared to the highest pork (0.01%) and beef (0.00%) content among the other 28 HPP samples (Supplementary Data by Haiminen *et al*.^38^). The microbiomes of these matrix contaminated samples also clustered into a separate sub-cluster (Figure 2B). This demonstrated that a shift in the food matrix composition was associated with an observable shift in the food microbiome.

MFMB-04 and MFMB-20 had the highest percentage of microbial reads compared to other samples (Supplementary Figure S3). They also exhibited an increase in *Lactococcus, Lactobacillus*, and *Streptococcus* relative abundances compared to other samples (Figure 5B), also reflected at respective higher taxonomic levels above genus (Supplementary Figure S5).

There were 53 genera identified uniquely in MFMB-04 and/or MFMB-20, but not present in any other sample. (MFMB-38 had a very low microbial load and contributed no uniquely identified genera above the abundance threshold.) MFMB-04 contained 44 unique genera (Figure 4) with the most abundant being *Macrococcus* (35.8 RPM), *Psychrobacter* (23.8 RPM), and *Brevibacterium* (18.1 RPM). Additionally, *Paenalcaligenes* was present only in MFMB-04 and MFMB-20 with an RPM of 6.4 and 0.3, respectively, compared to a median RPM of 0.004 among other samples. Notable differences in the matrix-contaminated samples’ unique microbial community membership compared to other samples may provide microbial indicators associated with unanticipated pork or beef presence.

### 2.6. Genus level identification of foodborne microbes

We evaluated the ability of total RNA sequencing to identify genera of commonly known foodborne pathogens within the microbiome. We focused on fourteen pathogen-containing genera including *Aeromonas, Bacillus, Campylobacter, Clostridium, Corynebacterium, Cronobacter, Escherichia, Helicobacter, Listeria, Salmonella, Shigella, Staphylococcus, Vibrio*, and *Yersinia* that were found to be present in the HPP samples with varying relative abundances. Of these genera, *Aeromonas, Bacillus, Campylobacter, Clostridium, Corynebacterium, Escherichia, Salmonella*, and *Staphylococcus* were detected in every HPP with median abundance values between 0.58–48.31 RPM (Figure 6A). This indicated that a baseline fraction of reads can be attributed to foodborne microbes when using NGS. Of those genera appearing in all samples, there was observed sample-to-sample variation in their abundance with some genera exhibiting longer tails of high abundance, e.g. *Staphylococcus* and *Salmonella*, whereas others exhibit very low abundance barely above the threshold of detection, e.g. *Bacillus* and *Yersinia* (Figure 6A). None of the pathogen-containing genera were consistent with higher relative abundances due to differences in food matrix composition. *Bacillus* and *Corynebacterium* exhibited slightly higher relative abundances in sample MFMB-04 which contained 7.7% pork and 9.0% beef (Figure 6B). Yet while MFMB-04 contained higher cumulative levels of these foodborne microbes, the next highest sample was MFMB-93 which was not associated with altered matrix composition, and both MFMB-04 and MFMB-93 contained higher levels of *Staphylococcus* (Figure 6B). Thus, matrix composition alone did not explain variations of these pathogen-containing genera.

**Figure 6A:**
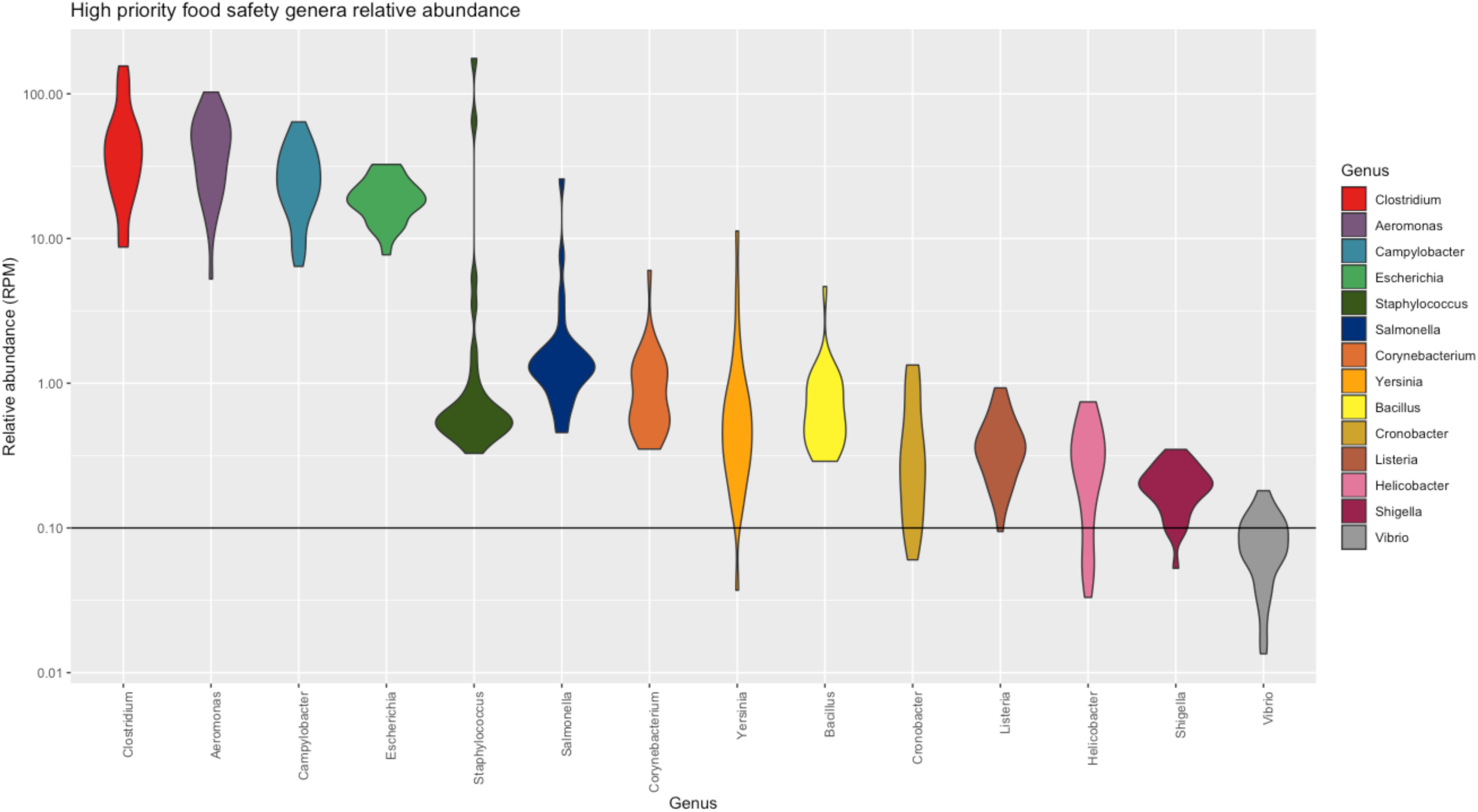
Relative abundance of microbes with high relevance to food safety and quality from high protein powder total RNA sequenced microbiomes. Width of violin plot indicates density of samples with relative abundance at that value. Observation threshold of RPM = 0.1 is indicated with the horizontal black line.

**Figure 6B:**
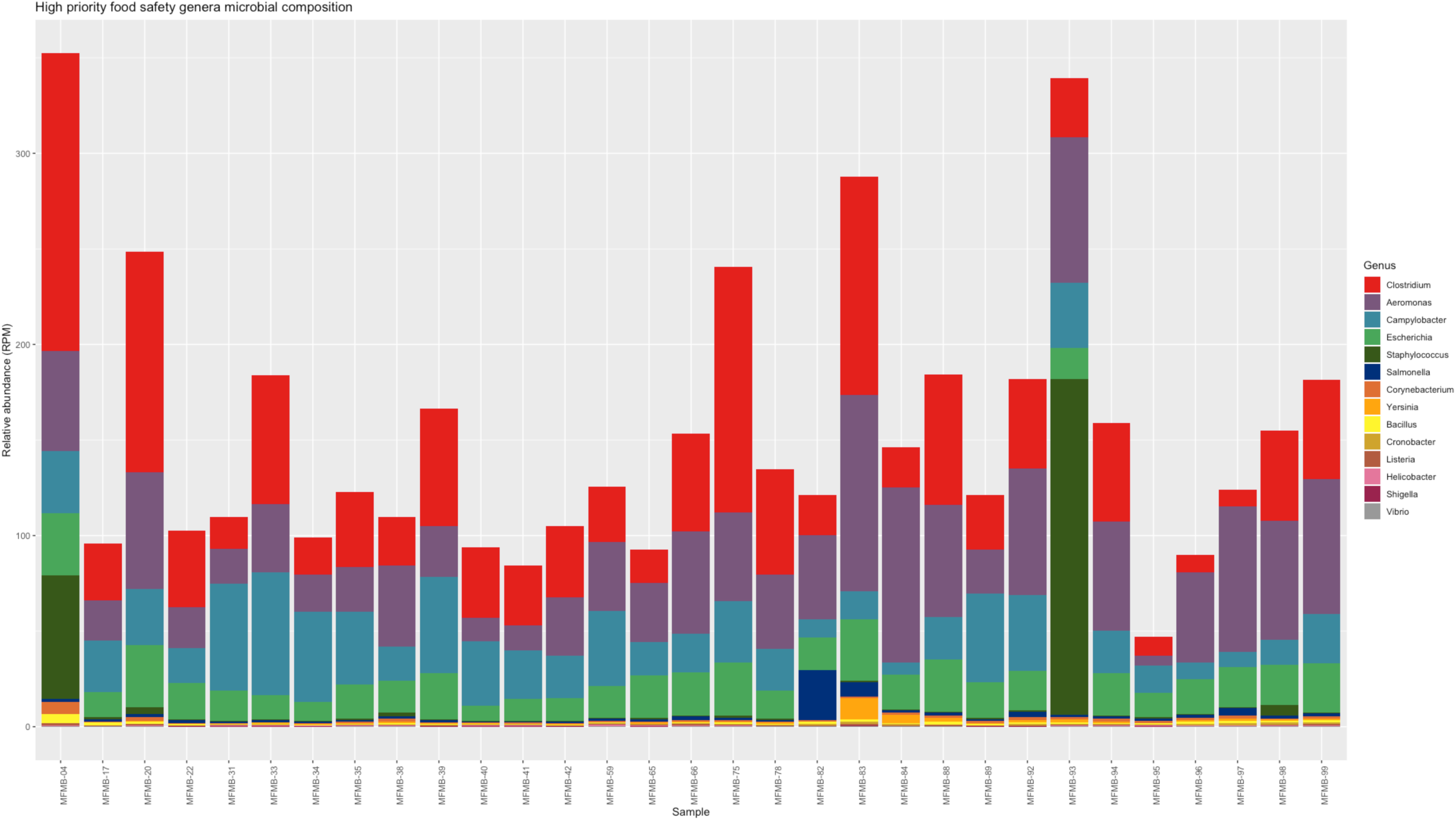
Foodborne microbe relative abundances are shown across samples of high protein powder total RNA sequenced samples.

Interestingly, low to moderate levels of *Salmonella* were detected within all 31 HPP microbiomes (Figure 6A). The presence of *Salmonella* in HPP is expected but the viability of *Salmonella* is an important indicator of safety and quality. Thus, we further sought to delineate *Salmonella* growth capability within these microbiomes by comparing culturability with multiple established bioinformatic NGS methods for *Salmonella* relative abundances in the samples.

### 2.7 Assessment of *Salmonella* culturability and total RNA sequencing

Total RNA sequencing of food microbiomes has the potential to provide additional sensitivity beyond standard culture-based food safety testing to confirm or reject the presence of potentially pathogenic microbes. In all of the examined HPP samples, some portion of the sequenced reads were classified as belonging to pathogen-containing genera (Figure 6); however, the presence of RNA transcripts does not necessarily indicate current growth of the organism itself. We further inspected one pathogen of interest, *Salmonella*, to determine the congruence between sequencing-based and culturability results. Of the 31 samples examined with total RNA sequencing, *Salmonella* culture testing was applied to 27 samples, of which four were culture-positive. Surprisingly, *Salmonella* culture-positive samples were not among those with the highest relative abundance of *Salmonella* from sequencing (Figure 7A). When ranking the samples by decreasing *Salmonella* abundance, the culture-positive samples were not enriched for higher ranks (*p*=0.86 from Wilcoxon rank sum test indicating that the distributions are not significantly different, Table 2). To confirm that the microbiome analysis pipeline did not miss *Salmonella* reads present, we completed two orthogonal analyses on the same data set used in the microbial identification step. The reference genomes relevant to these additional analyses were publicly available and closed high quality genomes available from the sources indicated below.

**Table 2:** The ranks for *Salmonella*-positive samples and the associated p-values from Wilcoxon rank sum test are shown for high-throughput sequencing read abundance (RPM) for multiple analyses: *k*-mer classification to NCBI Microbial RefSeq Complete (left), alignments to 1,447 *Salmonella* genomes (middle), and alignments to 4,846 *ef-Tu* gene sequences (right). The corresponding *Salmonella* relative abundances are shown in Figure 7A–C.

| *Salmonella*-positive sample | k-mer Classification | Whole Genome Alignment | *ef-Tu* Alignment |
| --- | --- | --- | --- |
| MFMB-04 | 8th | 10th | 1st |
| MFMB-20 | 9th | 9th | 4th |
| MFMB-38 | 20th | 3rd | 21st |
| MFMB-41 | 30th | 6th | 28th |
| Rank sum test p-value | p=0.86 | p=0.06 | p=0.56 |

**Figure 7:**
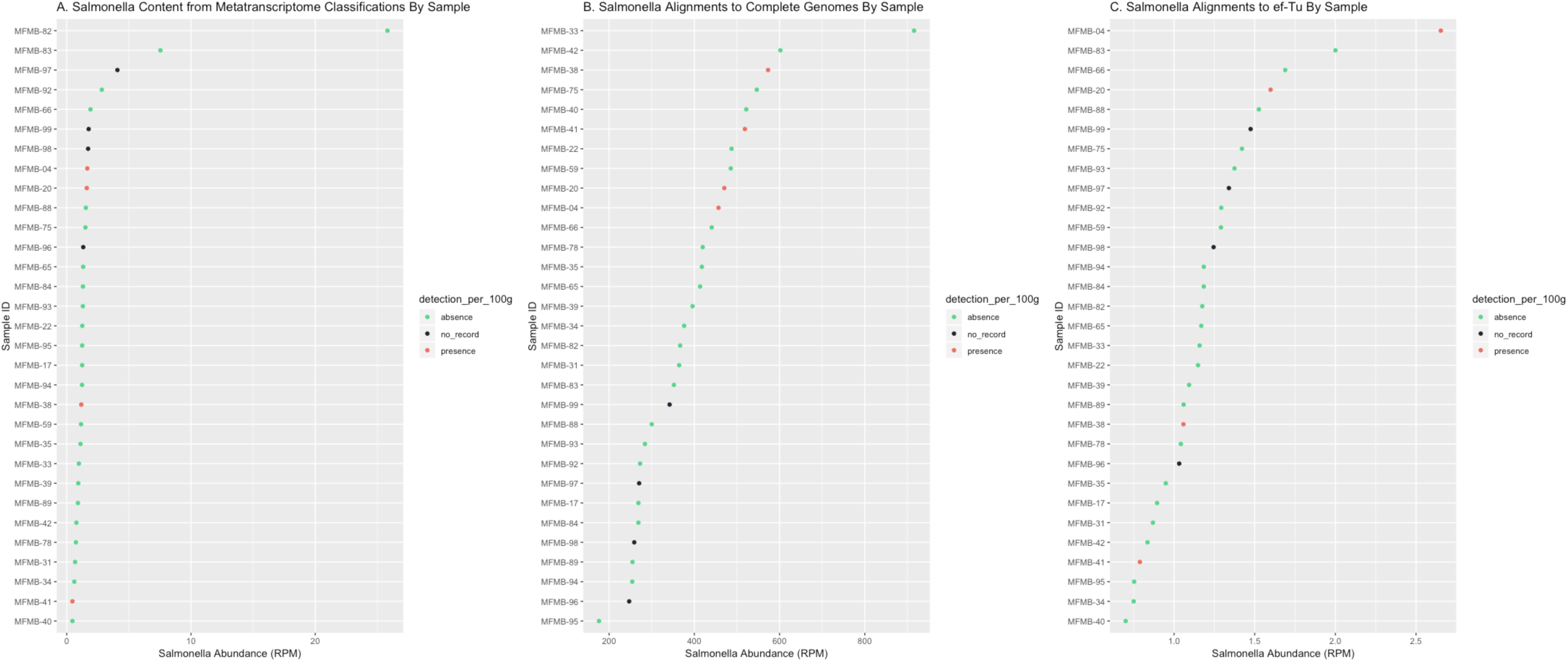
*Salmonella* culturability status and high-throughput sequencing read abundance (RPM) from *k*-mer classification to NCBI Microbial RefSeq Complete **(A)**, from alignments to 1,447 *Salmonella* genomes **(B)**, and from alignments to 4,846 EF-Tu gene sequences **(C)**. *Salmonella* presence (red) indicates culture-positive result, absence (green) indicates culture-negative result, and no record (black) indicates samples for which no culture test was completed.

First, for a targeted analysis, we aligned the sequenced reads using a different tool, Bowtie 2,^48^ to an augmented *Salmonella*-only reference database. This reference was comprised of the 264 *Salmonella* genomes extracted from NCBI RefSeq Complete (used in our previous microbial identification step) as well as an additional 1,183 public *Salmonella* genomes which represent global diversity within the genus.^49^ The number of reads that aligned to the *Salmonella*-only reference was on average 370-fold higher than identified as *Salmonella* by Kraken using the multi-microbe NCBI RefSeq Complete. In this additional analysis, the culture-positive samples had overall higher ranks compared to culture-negative samples (p=0.06, Table 2) indicating that additional *Salmonella* genomic data in the reference significantly improved discriminatory identification power. *Salmonella* culture-positive samples were still not the most abundant (Figure 7B), but with an enriched database, sequencing positioned all four culturable samples within the top 10 ranking.

The second additional analysis examined alignment of the reads to a specific gene required^50^ for replication and protein production in actively dividing *Salmonella—* elongation factor Tu (*ef-Tu*). This was done by aligning the reads to 4,846 gene sequences for *ef-Tu* extracted for a larger corpus of *Salmonella* genomes from OMXWare.^51^ The relative abundances of this transcript in culture-positive samples were still comparable to culture-negative samples (Figure 7C). Culture-positive samples did not exhibit higher ranks compared to culture-negative samples (p=0.56, Table 2), indicating that *ef-Tu* relative abundance alone was not sufficient to improve the lack of concordance in culturability vs sequencing. These two orthogonal analyses demonstrated that results from carefully developed culture-based testing and those from current high-throughput sequencing technologies, whether assessed at overall reads aligned or specific gene abundances, were not conclusively in agreement when detecting active *Salmonella* in food samples (Figure 7 and Table 2). However, the use of a reference database enriched in whole genome sequences of the specific organism of interested was found appropriate for food safety applications.

Since microbes compete for available resources within an environmental niche and therefore impact one another,^52^ we investigated *Salmonella* culture results in conjunction with co-occurrence patterns of other microbes in the total RNA sequencing data (Figure 8). Point-biserial correlation coefficients (*r*_*pb*_) were calculated between *Salmonella* culturability results (presence or absence which were available for 27 of the 31 samples) and microbiome relative abundance. We observed 31 genera that positively correlated and with *Salmonella* presence (*r*_*pb*_ > 0.5). *Erysipelothrix, Lactobacillus, Anaerococcus, Brachyspira*, and *Jeotgalibaca* exhibited the largest positive correlations. *Gyrovirus* was negatively correlated with *Salmonella* growth (*r*_*pb*_ = *−*0.54). In three of the four *Salmonella*-positive samples (MFMB-04, MFMB-20, and MFMB-38), food matrix contamination was also observed (Supplementary Data in Haiminen *et al*.^38^). The concurrency of *Salmonella* growth and matrix contamination was affirmed by the microbial co-occurrence (specifically *Erysipelothrix, Brachyspira*, and *Gyrovirus*). This highlights the complex dynamic and community co-dependency of food microbiomes, yet shows that multiple dimensions of the data (microbiome composition, culture-based methods, and microbial load) will signal anomalies from typical samples when there is an issue in the supply chain.

**Figure 8:**
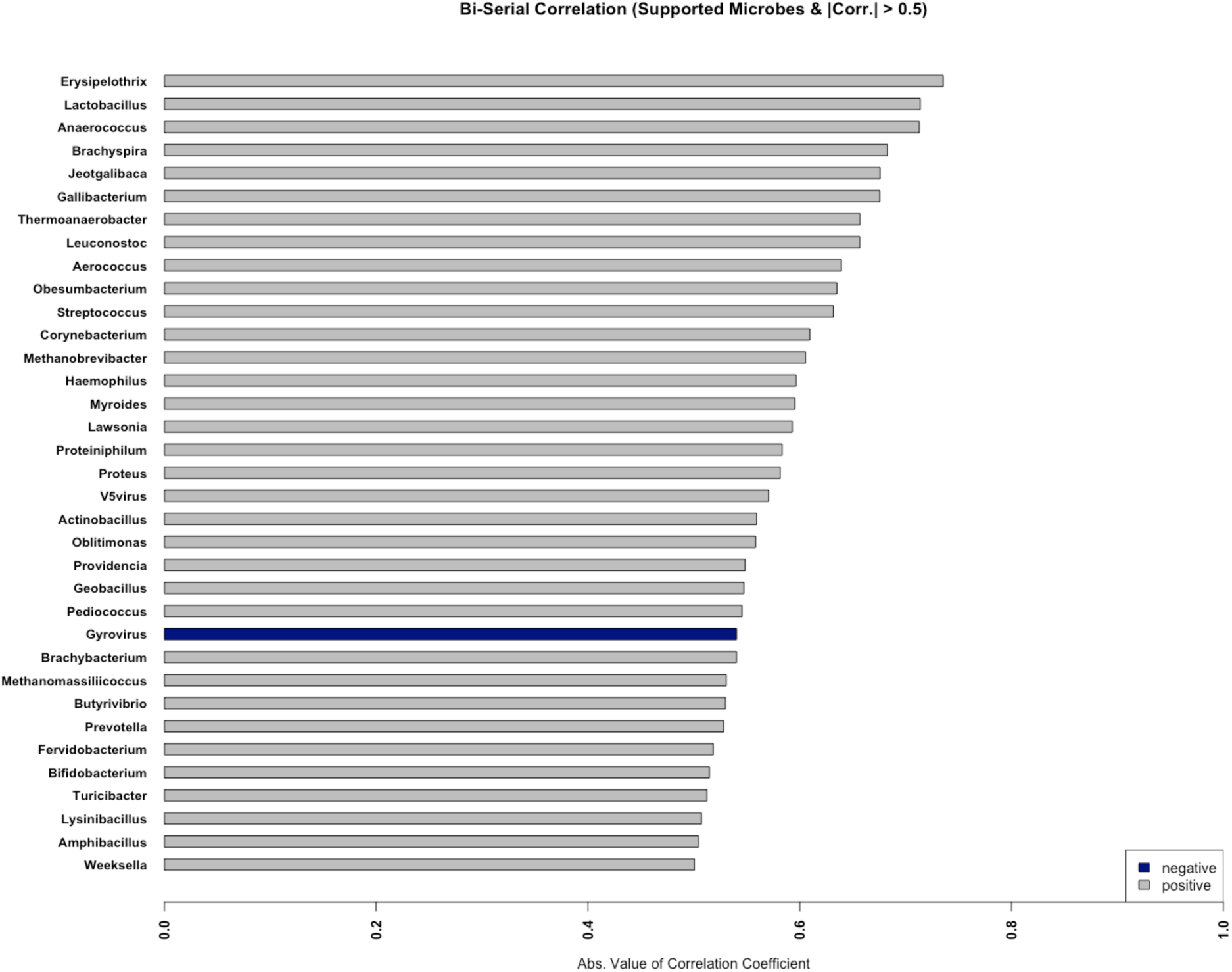
*Salmonella* status correlations with genus relative abundances. Only those genera with absolute value of the correlation coefficient > 0.5 are shown. Positive and negative correlations are indicated in grey and blue, respectively.

## 3. DISCUSSION

Accurate and appropriate tests for detecting potential hazards in the food supply chain are key to ensuring consumer safety and food quality. Monitoring and regular testing of raw ingredients can reveal fluctuations within the supply chain that may be an indicator of an ingredient’s quality or of a potential hazard. Such quality is assessed by standardized tests for chemical and microbial composition to meet legal requirements and specifications from government agencies throughout the world. For raw materials or finished products to meet these bounds of safety and quality, their composition must usually have a low microbiological load (except in fermented foods) and be chemically identical in macro-components such as carbohydrate, protein, and fat. Methods in this space must avoid false negative results which could endanger consumers, while also minimizing false positives which could lead to unnecessary recalls and food loss.

Existing microbial detection technologies used in food safety today such as pulse field gel electrophoresis (PFGE) and whole genome sequencing (WGS) require microbial isolation. This provides biased outcomes as it removes microbes from their native environment where other biotic members also subsist, and selects microbes by culturability alone. Amplicon sequencing, while a low-cost alternative to metagenome or metatranscriptome sequencing for bacteria, also imparts PCR amplification bias and reduces detection sensitivity due to reliance on a single gene (16S ribosomal RNA).^14,53,54^ We therefore investigated the utility of total RNA sequencing of food microbiomes and demonstrated that from this single test, we are able to yield several pertinent results about food safety and quality.

For this evaluation, we developed a pipeline to characterize the microbiome of typical food ingredient samples and to detect potentially hazardous outliers. Special considerations for food samples were made as computational pipelines for human or other microbiome analyses are not sufficient for applications in food safety without modification. In food, the eukaryotic matrix needs to be confirmed, may be mixed, and, as we and others have shown, affects the identification accuracy of microbes that are present.^35,36^ By filtering food matrix sequence data properly, we avoid incorrect microbial identification and characterization of the microbiome^36^ while also increasing the computational efficiency for downstream processing. The addition of this filtering step in the pipeline removed approximately 90% of false positive genera and provided results at 99.96% specificity when evaluating simulated mixtures of food matrix and microbes (Table 1).

Through the analysis of 31 high protein powder total RNA sequencing samples, we demonstrated the pipeline’s ability to characterize food microbiomes and indicate outliers. In this sample collection, we identified a core catalog of 65 microbial genera found in all samples where *Bacteroides, Clostridium*, and *Lactococcus* were the most abundant (Supplementary Table 4). We also demonstrated that in these food microbiomes the overall diversity was 2-fold greater than the core microbe set. Fluctuations in the microbiome can indicate important differences between samples as observed here, as well as in the literature for grape berry^6^ and apple fruit microbiomes (pertaining to organic versus conventional farming)^7^ or indicate inherent variability between production batches or suppliers as observed here and during cheddar cheese manufacturing.^8^ Specifically, we observed a shift in the microbial composition (Figure 2B) and the microbial load (Supplementary Figure S3) in high protein powder samples (derived from poultry meal) where unexpected pork and beef were observed. Matrix-contaminated samples were marked by increased relative abundances of specific microbes including *Lactococcus, Lactobacillus*, and *Streptococcus* (Figure 5B). This work shows that the microbiome shifts with observed food matrix contamination from sources with similar macronutrient content and thus, the microbiome alone is a likely signal of compositional change in food.

Beyond shifts in the microbiome, we focused on a set of well-defined foodborne-pathogen containing genera and explored their relative abundances observed from total RNA sequencing. Of these genera, *Aeromonas, Bacillus, Campylobacter, Clostridium, Corynebacterium, Escherichia, Salmonella*, and *Staphylococcus* were detected in every HPP sample. This highlights that when using NGS there may be an observable baseline of sequences assigned to potentially pathogenic microbes. For this ingredient type, this result lends a range of normalcy of relative abundance generated by NGS. Further work is needed to establish a definitive and quantitative range of typical variation in samples of a particular food source and the degree of anomaly for a new sample or genus abundance. However, preliminary studies of this nature can inform the development of guidelines when working with increasingly sensitive shotgun metagenomic or metatranscriptomic analysis.

Furthermore, sequenced DNA or RNA alone does not imply microbial viability. Therefore, we investigated the relatedness of culture-based tests and total RNA sequencing for the pathogenic bacterium *Salmonella* in the high protein powder samples. As has been reported for human gut^55^ and deep sea^56^ microbiomes, we also did not dretect a correlation between *Salmonella* read abundance and culturability (Figure 7 and Table 2). Sequence reads matching *Salmonella* references were observed for all samples (both culture-positive and culture-negative) as determined by multiple analysis techniques: microbiome classification, alignment to *Salmonella* genomes, and targeted growth gene analysis. When ranking the high protein powder samples based on *Salmonella* abundance from whole genome alignments, the culture-positive samples were enriched for higher ranks (*p* = 0.06). However, the culture-positive samples were still intermixed in ranking with culture-negative samples. This indicated that there was no clear minimum threshold of sequence data as evidence for culturability and that this analysis alone is not predictive of pathogen growth. One possible reason for this is that the culture-positive variant of *Salmonella* is missing from existing reference data sets. Potentially, *Salmonella* attained a nonculturable state wherein it was detected by sequencing techniques yet remained nonculturable from the HPP sources. Successful isolation of total RNA and DNA and gene expression analysis from experimentally known nonculturable bacteria has been demonstrated by Ganesan *et al*. in multiple studies in other genera.^19,22^ Physiological state should thus be taken under consideration when benchmarking sequencing technologies in comparison with culture-based methods. Thus, total RNA sequencing of food samples may identify shifts that standard food testing does not, but the incongruity between sequencing read data and culture-based results highlights the need to perform more benchmarking in food microbiome analysis for pathogen detection.

The characterization of HPP food microbiomes leveraged current accepted public reference databases, yet it is known that these databases are still inadequate.^1,2,11,57,58^ Furthermore, when considering congruence between *Salmonella* culturability and NGS read mapping techniques, the genetic breadth and depth of multi-genome reference sequences is essential. For example, focusing on *ef-Tu*, a known marker gene for *Salmonella* growth, was not sufficient to mirror viability of *in vitro* culture tests. This highlights the limitations of single gene approaches for identification. When the sequenced reads were examined in the context of an augmented reference collection of *Salmonella* genomes, we observed improved ranking and read mapping rate for culture-positive samples (yet we did not achieve complete concordance). This improvement underlined the increased analytical robustness yielded from a multi-genome reference. We also recognize that the read mapping rate may be exaggerated as reads from non-*Salmonella* genomes could map to *Salmonella* in the absence of any other reference genomes. Overall for robust analysis and applicability to food safety and quality, microbial references must be expanded to include more genetically diverse representatives of pathogenic and spoilage organisms. Description of food microbiomes will only improve as additional public sequence data is collected and leveraged.

In our sample collection, 2-4% (effectively 5 to 14 million) of reads remain unclassified. The GC content distribution of unclassified reads matched microbial GC content distribution (Supplementary Figure S4) suggesting that these reads may have been derived from microbes missing from the current reference database that have not yet been isolated or sequenced. By sequencing the microbiome, we sampled environmental niches in their native state in a culture-independent manner and therefore collected data from diverse and potentially never-before seen microbes. Tracking unclassified reads will also be essential for monitoring food microbiomes. The inability to provide a name from existing references does not eliminate the possibility that the sequence is from an unwanted microbe or indicates a hazard. In addition to tracking known microbes, quantitative or qualitative shifts in the unclassified sequences might be used to detect when a sample is different from its peers.

We demonstrated the potential utility of analyzing food microbiomes for food safety using raw ingredients. This study resulted in the detection of shifts in the microbiome composition corresponding to unexpected matrix contaminants. This signifies that the microbiome is likely an important and effective hazard indicator in the food supply chain. While we have used total RNA sequencing for detection of microbiome membership, the technology has future applicability for detection of antimicrobial resistance, virulence, and biological function for multiple food sources, and for other sample types. Notably, while this pipeline was developed for food monitoring, with applicable modifications and identification of material-specific indicators, it can be applied to other microbiomes including human and environmental.

## 4. METHODS

### 4.1 Sample Collection, Preparation, and Sequencing

High protein powder (HPP, 2.5 kg) samples were each collected from a train car in Reno, NV, USA between April 2015 and February 2016 in four batches from two suppliers and shipped to the Weimer lab at the University of California, Davis (Davis, CA). Each HPP sample was composed of five sub-samples from random locations within the train car prior to shipment. Sample preparation, total RNA extraction and integrity confirmation, cDNA construction, and library construction for these samples was previously described by Haiminen et al.^38^

Sequencing was performed by BGI@UC Davis (Sacramento, CA) using Illumina HiSeq 4000 (San Diego, CA) with 150 paired end chemistry for each sample except the following: HiSeq 3000 with 150 paired end chemistry was used for MFMB-04 and MFMB-17. All total RNA sequencing data are available via the 100K Pathogen Genome Project BioProject (PRJNA186441) at NCBI (Supplementary Table 1).

For evaluation of total RNA sequencing for microbial classification in paired processing steps, total RNA and total DNA were extracted from the same sample and denoted as MFMB-03 and MFMB-08, respectively. Total RNA was extracted and sequenced as described above. Total DNA was extracted and sequenced as described previously.^10,59–64^ The Illumina HiSeq 2000 with 100 paired end chemistry was used for MFMB-03 and MFMB-08.

### 4.2 Sequence Data Quality Control

Illumina Universal adapters were removed and reads were trimmed using Trim Galore^65^ with a minimum read length parameter 50 bp. The resulting reads were filtered using Kraken^37^, as described below in Section 4.3, with a custom database built from the PhiX genome (NCBI Reference Sequence: NC_001422.1). Removal of PhiX content is suggested as it is a common contaminant in Illumina sequencing data.^66^ Trimmed non-PhiX reads were used in subsequent matrix filtering and microbial identification steps.

### 4.3 Matrix Filtering Process and Validation

Kraken^37^ with a *k*-mer size of 31 bp (optimal size described in the Kraken reference publication) was used to identify and remove reads that matched a pre-determined list of 31 common food matrix and potential contaminant eukaryotic genomes (Supplementary Table 2). These food matrix organisms were chosen based on preliminary eukaryotic read alignment experiments of the HPP samples as well as high-volume food components in the supply chain. Due to the large size of eukaryotic genomes in the custom Kraken^37^ database, a random *k*-mer reduction was applied to reduce the size of the database by 58% using kraken-build with option --max-db-size, in order to fit the database in 188 GB for in-memory processing. A conservative Kraken score threshold of 0.1 was applied to avoid filtering microbial reads. The matrix filtering database includes low complexity and repeat regions of eukaryotic genomes to capture all possible matrix reads. This filtering database with the score threshold was also used in the matrix filtering *in silico* testing as described below.

Matrix filtering was validated by constructing synthetic paired end reads (150 bp) using DWGSIM^67^ with mutations from reference sequences using the following parameters: base error rate (e) = 0.005, outer distance between the two ends of a read pair (d) = 500, rate of mutations (r) = 0.001, fraction of indels (R) = 0.15, probability an indel is extended (X) = 0.3. Reference sequences are detailed in Supplementary Table 3. We constructed two *in silico* mixtures of sequencing reads by randomly sampling reads from eukaryotic reference genomes. Simulated Food Mixture 1 was comprised of nine species with the following number of reads per genome: 2M cattle, 2M salmon, 1M goat, 1M lamb, 1M tilapia (transcriptome), 962K chicken (transcriptome), 10K duck, 1K horse, and 1K rat totaling 7.974M matrix reads. Simulated Food Mixture 2 contained 5M soybean, 4M rice, 3M potato, 2M corn, 200K rat, and 10K drain fly reads, totaling 14.210M matrix reads. Both simulated food mixtures included 1,000 microbial sequence reads generated from 15 different microbial species for a total of 15K sequence reads (Supplementary Table 3).

### 4.4 Microbial Identification

Remaining reads after quality control and matrix filtering were classified using Kraken^37^ against a microbial database with a *k*-mer size of 31 bp to determine the microbial composition within each sample. NCBI RefSeq Complete^68^ genomes were obtained for bacterial, archaeal, viral, and eukaryotic microorganisms (∼7,800 genomes retrieved April 2017). Low complexity regions of the genomes were masked using Dustmasker^69^ with default parameters. A threshold of 0.05 was applied to the Kraken score in an effort to maximize the F-score of the result (as demonstrated in Kraken’s operating manual.^70^ Taxa-specific sequence reads were used to calculate a relative abundance in reads per million (RPM; Equation 1) where *R*_*T*_ represents the reads classified per microbial entity (e.g. the genus *Salmonella*) and *R*_*Q*_ represents the number of sequenced reads remaining after quality control (trimming and PhiX removal) for an individual sample, including any reads classified as eukaryotic:

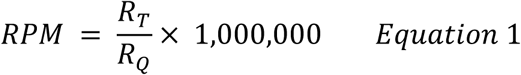

This value provides a relative abundance of the microbial entity of interest and was used in comparisons of taxa among samples. Genera with a conservative threshold of RPM > 0.1 were defined as present, as previously applied by others in the contexts of human infectious disease and gut microbiome studies.^33,34^ Pearson correlation of resulting microbial genus counts was computed.

### 4.5 Community Ecology Analysis

Rarefaction analysis at multiple subsampled read depths *R*_*D*_ was performed by multiplying the microbial genus read counts with R_D_/R_Q_ and rounding the results down to the nearest integer to represent observed read counts. Here R_Q_ is the total number of reads in the sample after quality control (including microbial, matrix, and unclassified reads). Resulting *α*-diversity at read depth R_D_ was computed as the number of genera with resulting RPM > 0.1 and plotted at five million read intervals: R_D_ = 5M, 10M, 15M, …, R_Q_. If, due to random sampling and rounding effects, the computed *α*-diversity was lower than the diversity computed at any previous depth, the previous higher *α*-diversity was used for plotting. The median elbow was calculated as previously described^45^ using the R package kneed.^45^

In compositional data analysis,^31^ non-zero values are required when computing *β*-diversity based on Aitchison distance.^46^ Therefore, reads counts assigned to each genus were pseudo-counted by adding one in advance of computation of RPM (Eq. 1) prior to calculating the Aitchison distance for the microbial table. *β*-diversity was calculated using the R package robCompositions^71^ and hierarchical clustering was performed using base R function hclust using the “ward.D2” method as recommended for compositional data analysis.^31^

### 4.6 Unclassified Read Analysis

The GC percent distributions of matrix (from matrix filtering), microbial, and remaining unclassified reads per sample were computed using FastQC^72^ and collated across samples with MultiQC.^73^

### 4.7 Analysis of *Salmonella* Culturability

Growth of *Salmonella* was determined using a real-time quantitative PCR method for the confirmation of *Salmonella* isolates for presumptive generic identification of foodborne *Salmonella*. Testing was performed fully in concordance with the Bacteriological Analytical Manual (BAM) for *Salmonella*^74,75^ for this approach that is also AOAC-approved. All samples with positive results for *Salmonella* were classified as containing actively growing *Salmonella*. To compare culture results with those from total RNA sequencing, *Salmonella* RPM values were parsed from the genus-level microbe table (described in Section 4.4).

Two additional approaches were employed to examine *Salmonella* read mapping with a more sensitive tool and broader reference databases. Quality controlled matrix-filtered reads were aligned using Bowtie2^48^ with very-sensitive-local-mode to 1. an expanded collection of whole *Salmonella* genomes and 2. to a curated growth gene reference for elongation factor Tu (*ef-Tu*). For results from both complete genome and *ef-Tu* gene alignments, the relative abundance (RPM) was computed as shown in Equation 1.

For whole genome alignments, a reference was constructed from 1,183 recently published *Salmonella* genomes^49^ in addition to the 264 *Salmonella* genomes extracted from the aforementioned NCBI RefSeq Complete collection (see Methods Section 4.4).

To construct a curated growth gene (*ef-Tu*) reference, gene sequences annotated in *Salmonella* genomes as “elongation factor Tu”, “EF-Tu” or “eftu” (case insensitive) were retrieved from the IBM Functional Genomics Platform^51^ (formerly known as OMXWare) using its Python package. This query yielded 4,846 unique gene sequences from a total of 36,242 *Salmonella* genomes which were assembled or retrieved from the NCBI Sequence Read Archive or RefSeq Complete Sequences as previously described.^51^ The retrieved *ef-Tu* gene sequences were subsequently used to build a custom Bowtie2^48^ reference. Read alignment was completed with very-sensitive-local-mode.

The read counts for each sample were ranked and Wilcoxon rank sum test was computed between the rank vectors of 4 *Salmonella*-positive and 23 *Salmonella*-negative samples. The 4 samples with unknown *Salmonella* status were excluded from the rankings.

Point-biserial correlation coefficients (*r*_*pb*_) were calculated between *Salmonella* growth indicated by culture results (+1 and -1 for presence and absence, respectively) and observed relative abundance from total RNA sequencing results using the R package ltm.^76^ The point-biserial correlation is a special case of the Pearson correlation that is better suited for a binary variable e.g. when *Salmonella* is reported as present or absent (a sample’s *Salmonella* status).

## Data Availability

All high protein powder (HPP) poultry meal sequences are available through the 100K Pathogen Genome Project (PRJNA186441) in the NCBI BioProject (see Supplementary Table 1 for a complete list of accession numbers).

## Code Availability

The pipeline and microbial or matrix references were constructed from publicly available tools and reference sequences as described in the Methods section. Automated usability of this pipeline is available through membership in the Consortium for Sequencing the Food Supply Chain.

## Acknowledgements

We’d like to acknowledge the IBM Research OMXWare team for their data management support and availability for the retrieval and processing of microbial genomes. This research project was financially supported by the Consortium for Sequencing the Food Supply Chain. Funding for the total RNA sequencing of high protein powder factory ingredients was provided by Mars, Incorporated to B.C.W. with specific interest in metagenomics of the food microbiome.

## Contributions

KLB and NH conceived of the experimental design, developed the approach, completed and oversaw the experiments, performed analyses, and wrote the paper; DC, SE, MK, BK, MD, RP, HK, ES developed the approach, analyzed data, and revised the manuscript; BCH completed nucleic acid extraction method development and sequencing library construction, and contributed to data analysis and writing; NK coordinated sample collection and processing, nucleic acid extraction and contributed to writing; RB and PM conceived of the experimental design, developed the approach, and reviewed the paper; BG contributed to the experimental design, developed the approach, and wrote the paper; GD, CHM, SP, AQ participated to the conception of the experimental design and to the review of the manuscript; LP conceived of the experiment, contributed to the data analysis, and wrote the paper; JHK conceived of the experiment, developed the approach, and wrote the paper; BCW conceived of the experimental design, developed the approach, oversaw the experiments, performed analyses, and wrote the paper

## Competing Interests

The authors were employed by private or academic organizations as described in the author affiliations at the time this work was completed. IBM Corporation, Mars Incorporated, and Bio-Rad Laboratories are members of the Consortium for Sequencing the Food Supply Chain. The authors declare no other competing interests

**Supplementary information is available at npj Science of Food’s website**

## SUPPLEMENTAL INFORMATION

Supplemental Figures (pdf): Supplemental Figures S1–S5

Supplemental Table 1 (.xlsx) - Sample Descriptions

Supplemental Table 2 (.xlsx) - Matrix Filtering Genomes

Supplemental Table 3 (.xlsx) - Simulated Food Mixtures

Supplemental Table 4 (.xlsx) - Microbial Genera

