## Supplemental Figures for "Monitoring the microbiome for food safety and quality using deep shotgun sequencing"

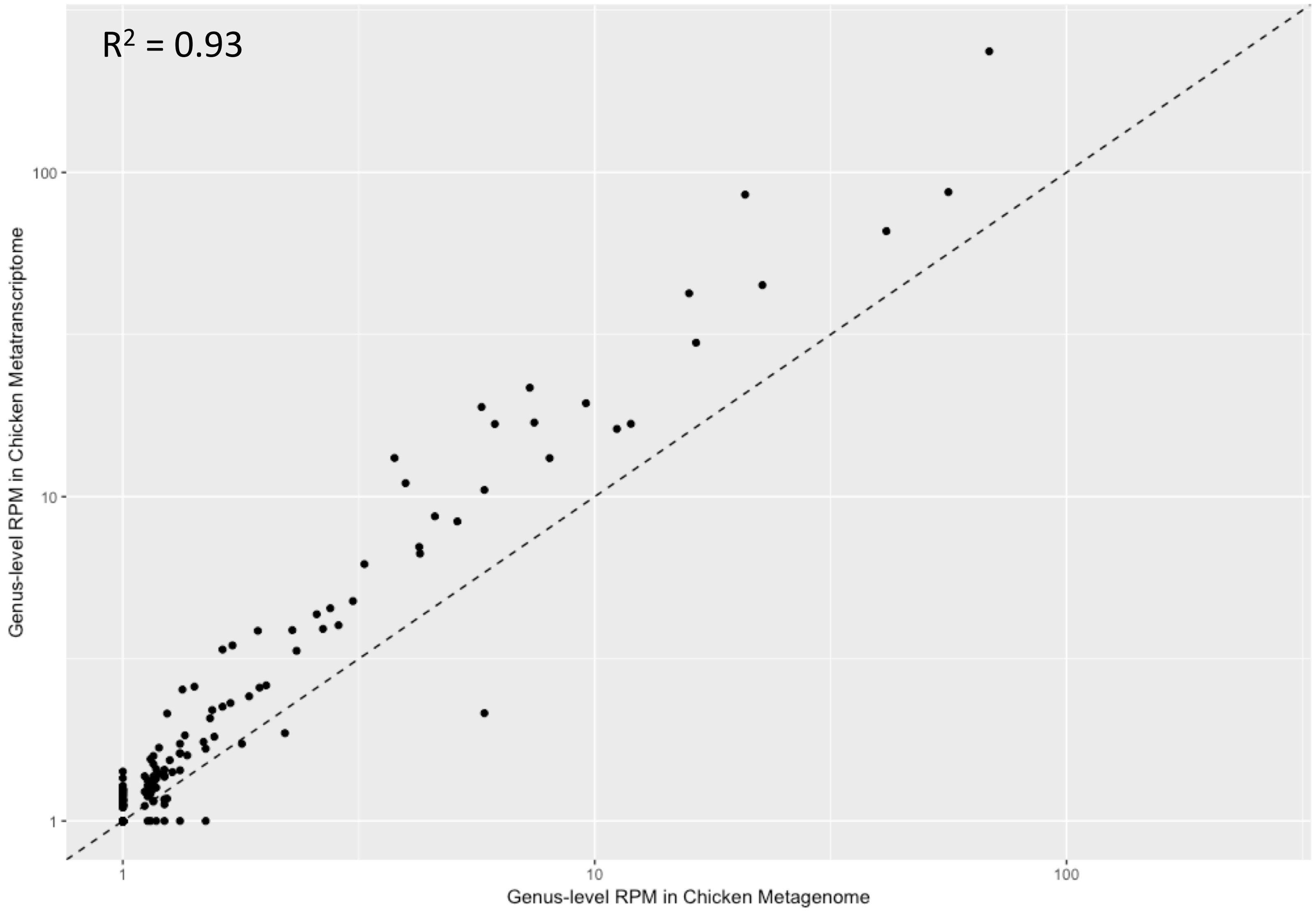


**Supplemental Fig. S1**: Genus-level relative abundance (RPM) in total RNA metatranscriptome and total DNA metagenome high protein powder samples of the same starting material.

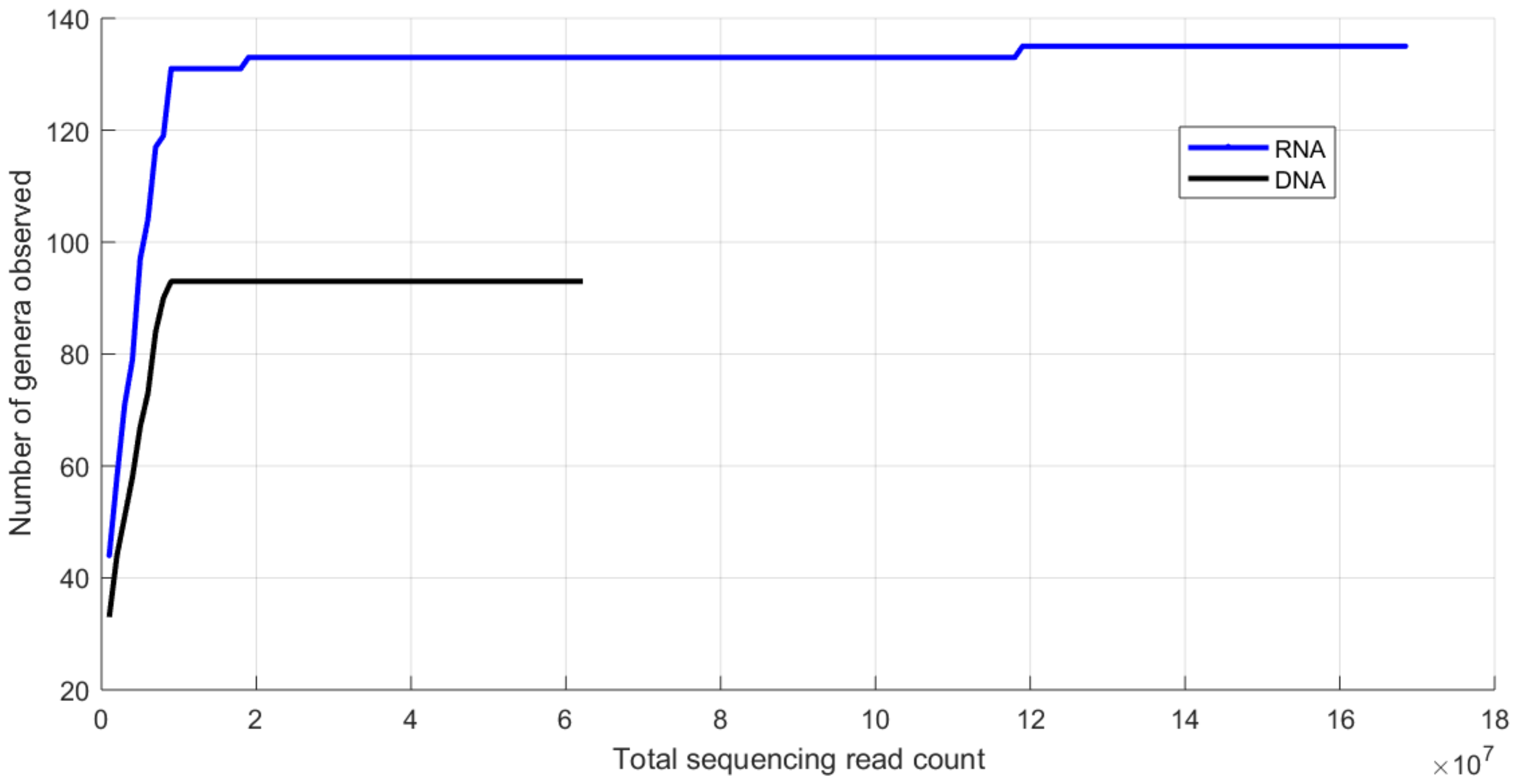


**Supplemental Fig. S2**: Alpha diversity (number of genera) for total RNA metatranscriptome and total DNA metagenome  sequencing of high protein powder samples of the same starting material, for a range of *in silico* subsampled sequencing depths (total sequence read counts including food matrix sequences).

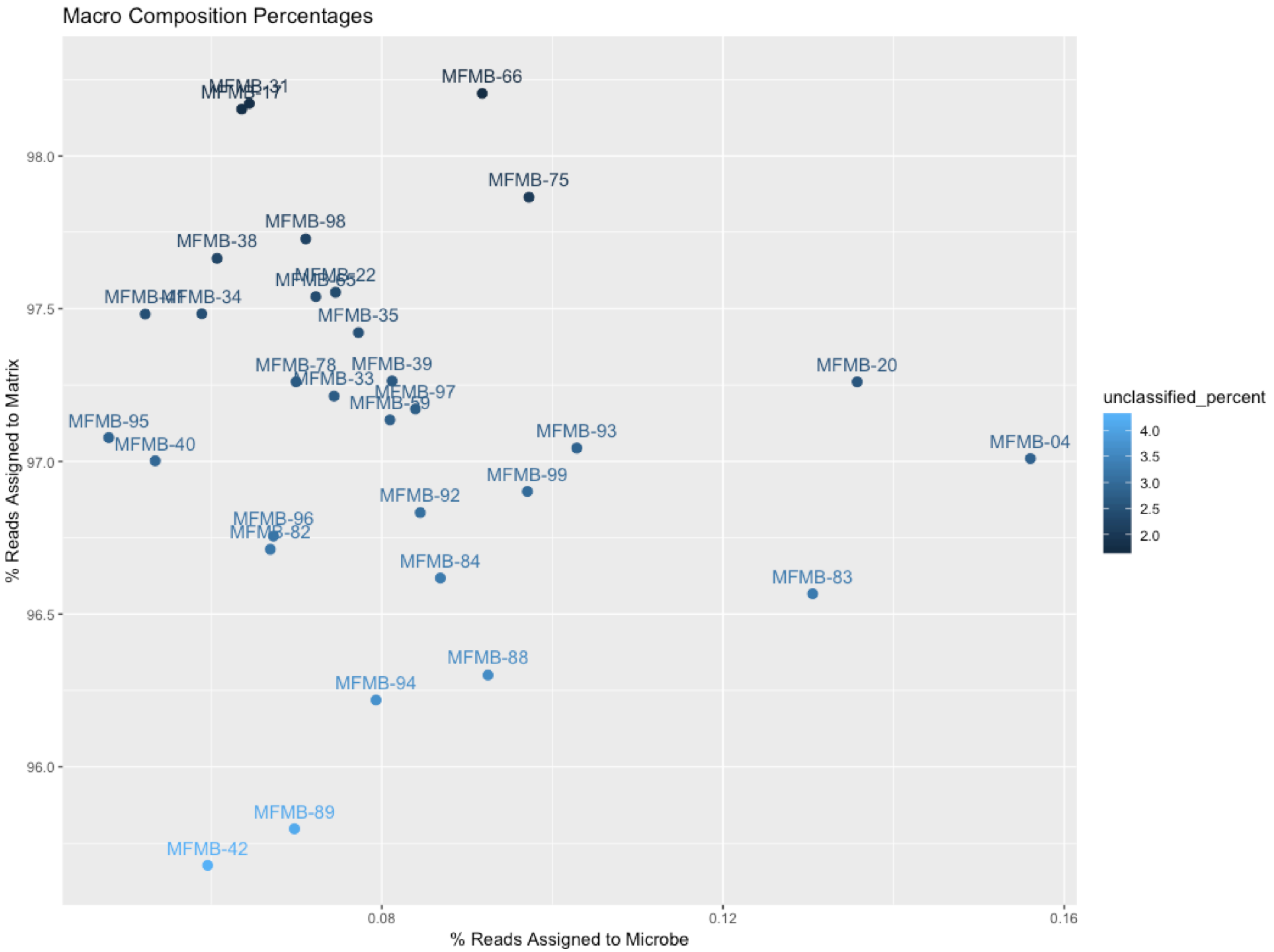


**Supplemental Fig. S3**: Macro-composition variations from total RNA metatranscriptome sequencing for each high protein powder sample (n=31): the percentage of reads passing quality control that were assigned to matrix, microbes, or could not be assigned (unclassified).

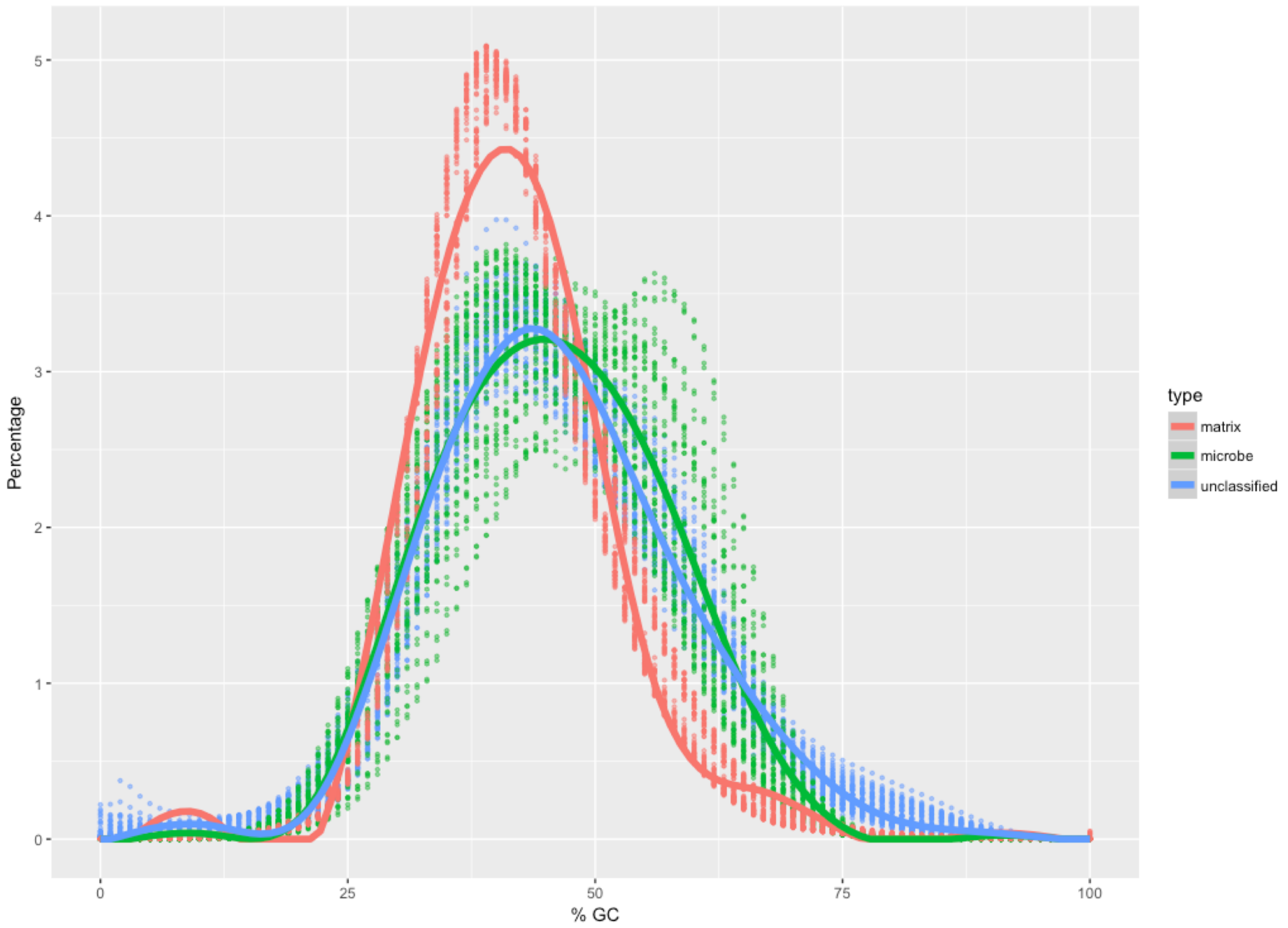


**Supplemental Fig. S4**: GC content of reads for three macro-composition origination groups: matrix (red), microbe (green) and unclassified (blue) for all samples (n=31) as points, with a trend line shown as a solid line using a generalized additive model.

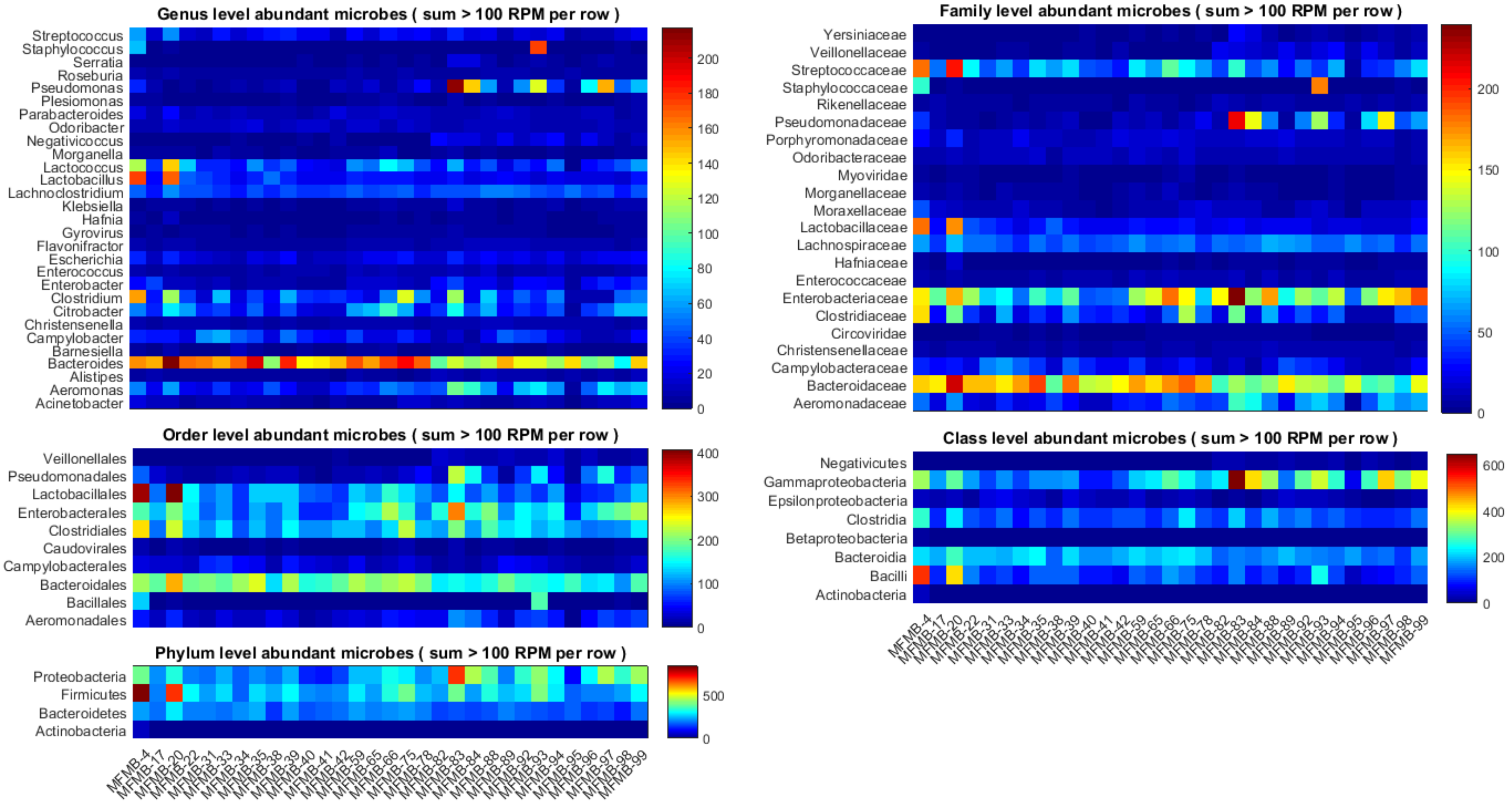


**Supplemental Fig. S5:** Most abundant microbes at various taxonomic levels, including only those with a sum of RPM > 100 across all samples.
